# SALL2 constrains TEAD4 by maintaining repressive chromatin to restrict trophectoderm identity

**DOI:** 10.64898/2026.07.10.737659

**Authors:** Yu Qiao, Jianfei Xu, Lin Zheng, Zhen Xiao, Zhuoqi Huang, Zhangting Liang, Xuemeng Zhou, Gang Ma, Guoqing Tong, Miguel A. Esteban, Andrew P. Hutchins

## Abstract

Embryonic development is marked by the successive restriction of developmental potential and the specification of embryonic and extraembryonic lineages. Yet, how these lineage decisions are established, and how the epigenome is remodelled to promote and restrict cell fate transitions, remains poorly understood. Here, we demonstrate that *SALL2* knockdown in primed human pluripotent stem cells (hPSCs) triggers a trophectoderm (TE)-like phenotype, characterized by palisade-like morphology and the up-regulation of TE-associated genes. Mechanistically, SALL2 physically interacts with the key TE driver TEAD4 and maintains bivalent, repressive chromatin (H3K4me3/H3K27me3) at TE-specific loci. Reduced *SALL2* led to enhanced TEAD4 occupancy and disrupted H3K27me3 and increased active chromatin marks at TE genes. Importantly, depletion of TEAD4 abolished the TE-like phenotype induced by *SALL2* knockdown, demonstrating that TEAD4 is required for the downstream effects of SALL2 loss. In support of this, blastoid-like aggregates can be generated from primed hPSCs with *SALL2* knocked down. Together, our findings identify SALL2 as a key epigenetic barrier that restrains TE lineage commitment by limiting TEAD4-dependent activation of the trophoblast transcriptional program.

**Key findings:**

- SALL2 acts as a molecular barrier to trophectoderm differentiation.
- SALL2 suppresses the trophectoderm by maintaining bivalent repressive chromatin at TEAD4-bound loci.
- Blastoid-like structures can be generated from primed-state media when *SALL2* was knocked down.

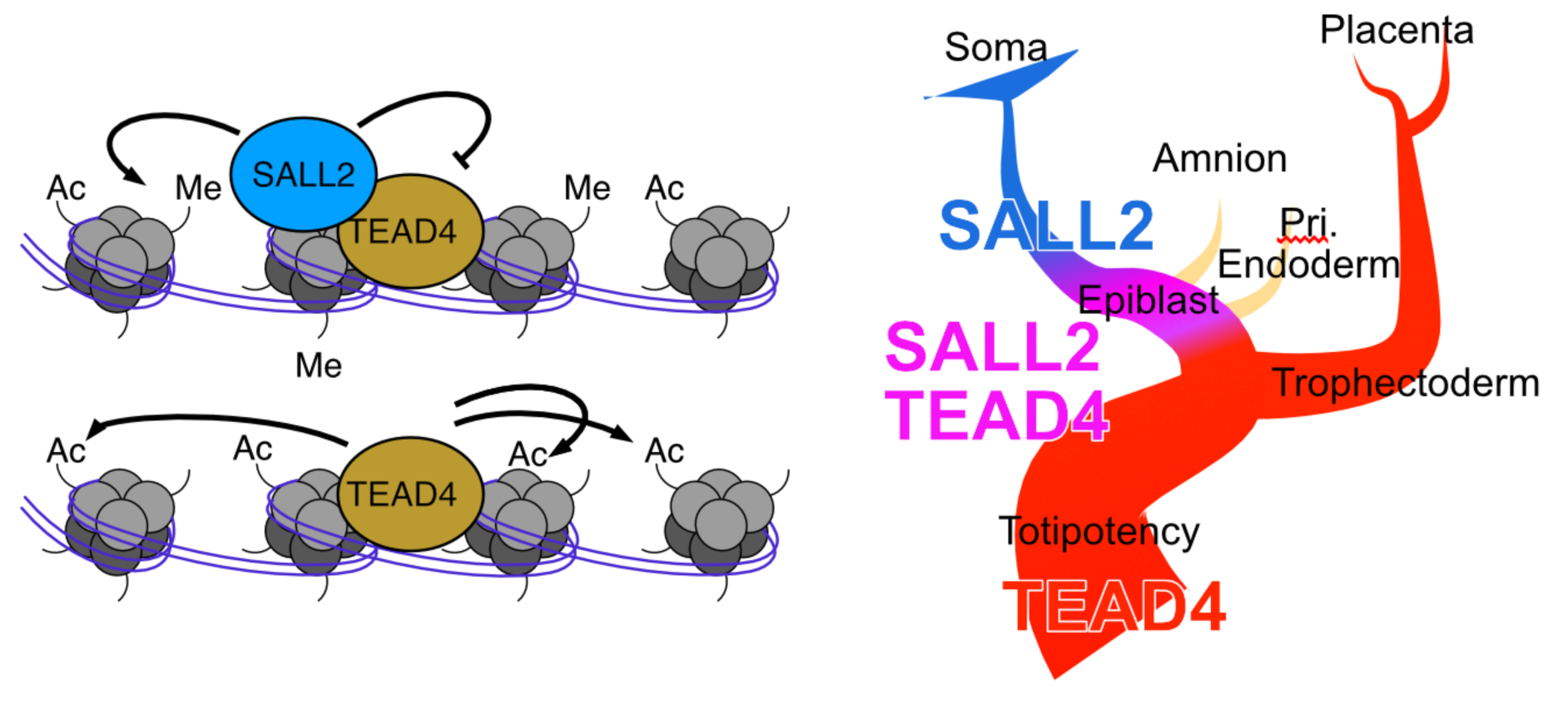

## Introduction

Early embryonic development is a very active time in cell transitions. One of the most important is the establishment of the trophectoderm (TE), which emerges from the external cells of the morula ^1–3^ to make up the placenta ^4^. How blastomeres transition from a totipotent state to the TE, and the inner cell mass (ICM) remains unclear, particularly in human ^5^. Understanding is especially hindered as mouse embryos use divergent mechanisms to specify ICM and TE compared to human and primate ^6–8^. In mouse, CDX2 competes with and represses OCT4 to specify TE just before the blastocyst stage ^8,9^. However, in humans, TEAD4/VGLL1/YAP and the parallel pathways based around GATA3/TFAP2C appear to be more important, and are upstream of CDX2 ^6,10–16^. Indeed, in human embryos CDX2 is detected much later than mouse, only after TE establishment at the blastocyst stage ^17^.

Here, we identified a role for the transcription factor SALL2 in TE development. When *SALL2* was knocked down in primed human pluripotent stem cells (hPSCs) they adopted characteristics of TE-like cells and TE-related genes were upregulated. Mechanistically we showed that SALL2 co-binds to TEAD4-bound loci to inhibit TEAD4’s ability to promote a TE cell fate. SALL2 inhibited TEAD4 by maintaining a bivalent chromatin state, that shifts to an activated chromatin state when SALL2 is removed. Strikingly, knockdown of *SALL2* resulted in improved blastoids in minimal conversion media. Our model suggests that, as *SALL2* is expressed after the split of the ICM and TE lineages, SALL2 acts as an epigenetic lock to stop reversion or transdifferentiation to a TE-fate and enable commitment to somatic lineages.

## Results

### *SALL2* knockdown induces TE gene expression in primed hPSCs

As an extension of our work to identify naïve and totipotent-related factors we previously identified transcription and epigenetic factors that are low in pre-blastocyst development and become high at later stages ^18^. In that analysis we also identified *SALL2* as low in pre-implantation development (**Supplementary Figure 1A, B**). SALL2 was also preferentially expressed in primed and differentiated hPSCs and was low in naïve hPSCs in a pattern similar to *HNRNPU* (**Supplementary Figure 1C-F**). Our expectation was that, like *HNRNPU*, knockdown of *SALL2* would cause cells to adopt a naïve or totipotent character. However, the cells instead formed flat cultures of palisade-like cells (**Figure 1A-C**). Pluripotent and naïve-specific genes (already low in primed hPSCs) were down-regulated in the *SALL2* KD cells (**Figure 1D**). The morphology of the cells resembled human trophectoderm stem cells (hTSCs) or trophectoderm-like cells (TELCs) (**Figure 1C**) ^10,11,19^, hence we performed RNA-seq to identify the cells. *SALL2* knockdown had a strong effect on the transcriptome, up-regulating 1233 genes and down-regulating 972 (**Figure 1E**). Comparison to a panel of RNA-seq data (See **Methods**) suggested the up-regulation of a hTSC/placental-gene expression program (**Figure 1F and Supplementary Figure 2A, B**). RNA-seq supported the up-regulation of hTSC-related genes and the RT-qPCR confirmed these changes (**Figure 1G, H**). Immunofluorescence detected cells expressing the TE-marker GATA3, which was mutually exclusive to cells expressing SOX2 when *SALL2* was knocked down (**Figure 1I**). Primed-specific genes were also downregulated (**Figure 1J**). Single cell RNA-seq supported the increased differentiation of sh*SALL2* cells by RNA velocity (**Figure 1K**), and the upregulation of TE-specific genes in *SALL2* knockdown cells (**Figure 1L, M**).

**Figure 1.**
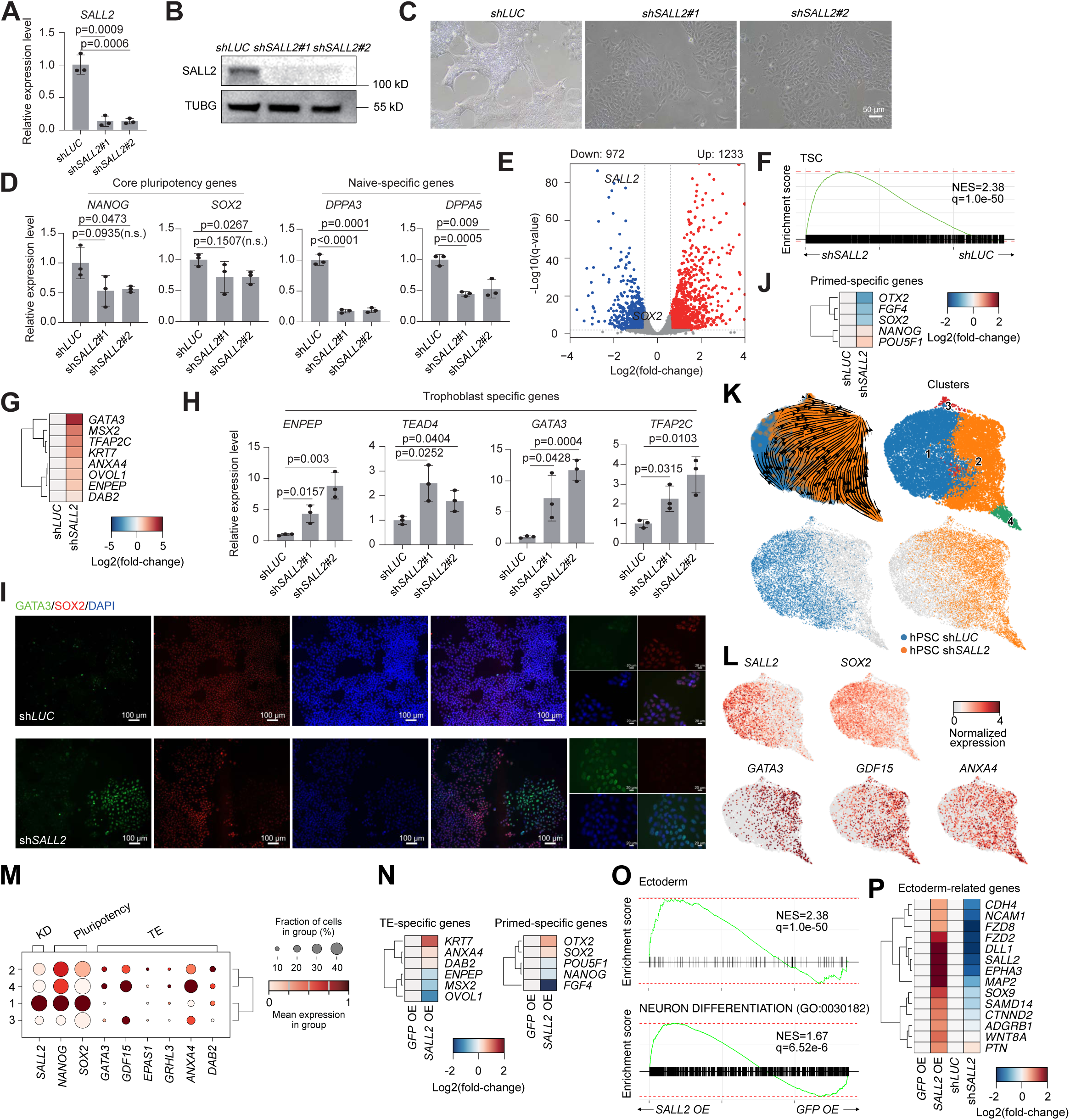
Knocking down *SALL2* in hPSCs upregulates TE-specific genes. **A** qRT-PCR showing the knockdown efficiency for *SALL2* in hPSCs. Target RNAs are normalized to *GAPDH*. The experiment was performed 3 times in biological replicate, each with three technical replicates, significance is from a two-tailed Welch’s t-test. **B** Western blot of SALL2 and TUBG in hPSCs transfected with an shRNA targeting luciferase as a control (sh*LUC*), or shRNAs targeting *SALL2*. The experiment was repeated twice with similar results. **C** Bright-field microscopy of hPSCs transfected with a control shRNA (shLUC) or with an shRNA targeting *SALL2* (sh*SALL2*#1, #2). Scale bar = 50 µm. The experiment was repeated three times with similar results. **D** RT-qPCR of the pluripotent genes *NANOG* and *SOX2* and the naïve-specific genes *DPPA3* and *DPPA5*. The experiment was performed in biological triplicate. **E** Volcano plot of RNA-seq for *SALL2* KD hPSCs. The experiment was performed in biological duplicate. Significance is from DESeq2 BH-corrected p-values. **F** GSEA forhTSC-specific genes. NES = normalized enrichment score; q = Bonferroni–Hochberg-corrected p-value. **G** Heatmap of the expression of selected TE-related genes. **H** RT-qPCR analysis of trophoblast specific genes (*ENPEP*, *TEAD4*, *GATA3* and *TFAP2C*). **I** Immunofluorescence staining of GATA3, and SOX2 at day 2 after puromycin selection in *SALL2* KD hPSCs. Scale bar = 100 µm. **J** Heatmap of the expression of selected primed-specific genes in the *SALL2* KD cells. **K** UMAP of scRNA-seq of hPSCs with knockdown of SALL2. **L** UMAPs of the selected expression of *SALL2*, the pluripotency gene *SOX2*, or the TE-related genes (*GATA3*, *GDF15*, and *ANXA4*). **M** Bubble plot showing the expression of selected pluripotency or TE-related genes. **N** Heatmap of selected TE and primed-specific genes in the SALL2 overexpressing cells. **O** GSEA of the up-regulated genes in hPSCs overexpressing *SALL2*. **P** Heatmap of selected ectoderm-related genes in hPSCs overexpressing *SALL2* or transfected with shRNAs targeting *LUC* or *SALL2*. Fold-change is relative to their respective controls.

Reduced *SALL2* biases cells to adopt a TE-like phenotype. Conversely, its overexpression not only reversed this trend but also further suppressed TE genes and boosted pluripotency markers (**Figure 1N**). Overexpression of *SALL2* did not induce a naïve phenotype (**Supplementary Figure 2C, D**), it did push the cells to express ectoderm-related genes (**Figure 1O, P**), in agreement with the reported roles for SALL2 in brain development and glioblastoma ^20,21^. Overall, these results demonstrate that depletion of SALL2 is sufficient to drive a cell-fate transition from primed pluripotency toward a trophoblast-like identity.

### Knockdown of *SALL2* improves the TE differentiation

Knockdown of *SALL2* up-regulates TE-specific genes, but the above experiments were performed in normal primed hPSC medium that lacks signaling cues for hTSC differentiation. Hence, we explored the impact of reduced *SALL2* on hTSC differentiation. hPSCs were differentiated to hTSCs using an established culture medium based on exogenous FGF ^19^. As expected, cell morphology changed to form flat palisade-like cells distinctive of hTSCs (**Figure 2A**), which were positive for the hTSC-markers KRT7 and GATA3 (**Figure 2B**). RNA-seq of the differentiation time course showed broad agreement between the *LUC* and *SALL2* knockdowns (**Figure 2C**), although PCA indicated some divergence in the differentiation trajectory along PC4 (**Figure 2D**), and each time point showed several hundred gene expression changes (**Supplementary Figure 3A**). GSEA of significantly differentially regulated genes for sh*SALL2* versus sh*LUC* on day 6 suggested that the *SALL2* knockdown hTSC-like cells were differentiating towards a placental-like cell fate more efficiently (**Figure 2E**). Indeed, specific hTSC-marker gene expression was accelerated in the *SALL2* KD cells, including *CDX2*, *ANXA4*, *CGA* and *DAB2* (**Figure 2E, F**). Pluripotency genes, however, tended to be unchanged, and genes representing the epithelial-mesenchymal transition (EMT) that accompanies hTSC differentiation were also mostly unaffected (**Figure 2E**). Together, these data suggests that *SALL2* KD accelerates the emergence of hTSC gene expression but does not alter the pluripotency suppression nor alter the EMT.

**Figure 2.**
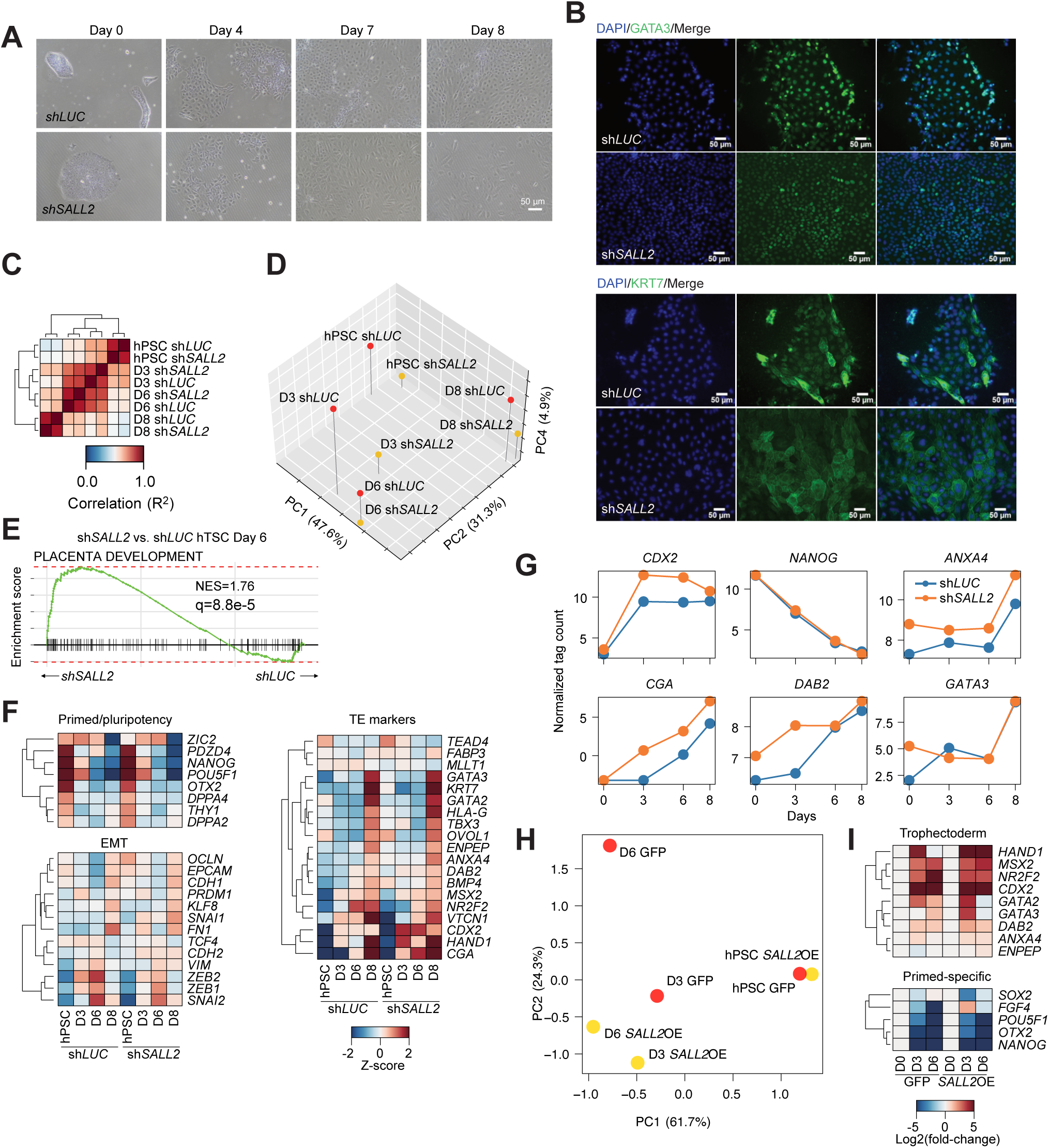
*SALL2* knockdown promotes the differentiation of hPSCs to hTSCs. **A** Morphological changes with days in H1 SALL2 KD hESCs and H1 after hTSC induction at days (D) 0, 4, 7 and 8. Scale bar = 50 μm. **B** Immunofluorescent staining of KRT7, and GATA3 at day 7 after hTSC induction. **C** Cross-correlation of all pair-wise RNA-seq samples for the differentiation of hPSCs to hTSCs at the indicated days. **D** PCA of the RNA-seq of the differentiation time course of hPSCs differentiated to hTSCs transfected with shRNAs targeting *SALL2* or *LUC* as a control. **E** GSEA of genes at day 6 of the hTSC differentiation. **F** Heatmap of selected pluripotent, epithelial-mesenchymal transition (EMT) genes, or TE-related genes over the hTSC course time in cells transfected with a control sh*LUC* or with an shRNA targeting *SALL2*. **G** Selected line plots showing TE-related changes over time in hTSC differentiation time course. **H** PCA of the RNA-seq of the differentiation time course of hPSCs differentiated to hTSCs transfected overexpressing *SALL2* or *GFP* as a control. **I** Heatmap of selected TE or hPSC primed-specific genes in a hTSC differentiation time course in cells overexpressing *SALL2* of *GFP* as a control

*SALL2* knockdown promoted hTSC differentiation, hence we predicted *SALL2* overexpression would have the opposite effect. As expected, although the overall trajectory towards a TE fate was unaffected by the overexpression of *SALL2*, cross-correlation and PCA suggested the day 6 *SALL2* overexpressing cells were closer to the day 3 GFP controls (**Supplementary Figure 3B, C**). This was supported by the increased number of significantly differentially regulated genes at day 6 (**Supplementary Figure 3D**). However, TE marker genes were upregulated mostly as usual, and pluripotency downregulated as expected (**Figure 2H, I and Supplementary Figure 3E, F**). These data indicate that exogenous signaling factors can overcome high *SALL2* expression. However, overall, combined with the knockdown data for *SALL2* it suggests SALL2 primarily acts to restrain TE differentiation.

Whilst hTSCs can be formed from both naïve and primed cells ^22,23^, it is more efficient when the starting cells are in a naïve-like state ^24–26^. Reanalysis of primed versus naïve (EPSC) differentiation shows that *SALL2* is lower in the naïve EPSC cells compared to primed (**Supplementary Figure 4A**) ^26^. This was also observed in PGXL naïve cells differentiated to hTSCs (**Supplementary Figure 4B, C**) ^24^. To confirm this, we reanalyzed a scRNA-seq data set for hTSC differentiation. From both naïve and primed hPSCs and embedded our hPSC knockdown *SALL2* and *LUC* cells (**Supplementary Figure 4D, E**) ^25^. As in the PGXL naïve cells, SALL2 was also lower expressed in the 5iLAF naïve cells in this protocol (**Supplementary Figure 4F, G**). Interestingly, the hTSC differentiation from naïve cells did not alter the direction of differentiation, instead it promoted hTSC emergence, as shown by the increased number of Naïve day 6 cells in clusters 4 and 6 and in the final hTSCs (**Supplementary Figure 4H**). We embedded our sh*SALL2* knockdown cells into this scRNA-seq data set and found that a small percentage (∼3%) of the *SALL2* knockdown cells clustered with hTSCs (**Supplementary Figure 4I**). It should be emphasized that the *SALL2* knockdown cells were grown in primed hPSC media and there is no exogenous activation of hTSC signaling pathways. These data suggest that *SALL2* is a barrier to hTSC differentiation, and at least one of the mechanisms by which naïve cells show improved hTSC differentiation is due to their lower level of SALL2. Indeed, in an RNA-seq panel of hPSC-naïve culture conditions *SALL2* is uniformly lower (**Supplementary Figure 4J, K**). This data suggests low *SALL2* is important for priming naïve cells for efficient hTSC differentiation.

### SALL2 and TEAD4 co-bind to genomic loci to regulate TE-specific genes

We next set out to explore how SALL2 inhibits TE differentiation. CUT&Tag for SALL2 revealed 91,065 bound loci. SALL proteins do not have a well-defined DNA binding sequence motif, as they bind to AT-rich regions ^27^. Motif discovery did not identify a strong motif for SALL2 but did identify several potential co-factors (**Figure 3A**). We downloaded genome-wide binding for these predicted co-factors, and the data suggested SALL2 was indeed co-binding to the genome along with JUN, MAX, and POU5F1 (OCT4) (**Supplementary Figure 5A, B**). One particularly interesting co-factor was TEAD4, which is a driver of trophoblast development ^10,16,28^, as overexpression in mouse causes differentiation to a TE-like fate ^13,14^, and is essential for hTSC self-renewal ^29,30^. Interestingly, TEAD4 is expressed at high levels in both naïve and primed hPSCs (**Supplementary Figure 4J**). Reanalysis of TEAD4 binding showed substantial overlap between the two factors (**Figure 3B**). We confirmed a direct interaction between SALL2 and TEAD4 using co-IP/Western. Interestingly, whilst SALL2 can precipitate TEAD4, TEAD4 did not precipitate SALL2 (**Figure 3C**), suggesting the physical interaction is tenuous. We performed Co-IP/MS with antibodies against both SALL2 and TEAD4 (**Supplementary Figure 5C and Supplementary Table 1**). SALL2 and TEAD4 were identified in the overlapping Co-IP/MS (**Figure 3C, Supplementary Figure 5C and Supplementary Table 1**). Of the potential transcription factors identified in **Supplementary Figure 5C**, only POU5F1 and TEAD4 were precipitated, suggesting a weak or indirect interaction with those other factors. Known co-factors for TEAD4 include p53 ^31^, which did not co-precipitate with TEAD4, however, it was bound to the genome at similar loci to SALL2 and TEAD4 (**Supplementary Figure 5D**). Interestingly, p53 was reduced at SALL2/TEAD4 bound loci in hTSCs (**Supplementary Figure 5D**). Conversely, other TEAD4 co-factors SP6 and VGLL1 ^10,11,32^ are not expressed in hPSCs, but reanalysis of C&T data for VGLL1 in hTSCs and TELCs (day 5) and ChIP-seq for SP6 in cytotrophoblast-like cells shows that SALL2 bound-loci are sites of future recruitment for both VGLL1 and SP6, but not p53 in TE (**Supplementary Figure 5D**). These data show that SALL2 is bound to TE-related regulatory elements in primed hPSCs.

**Figure 3.**
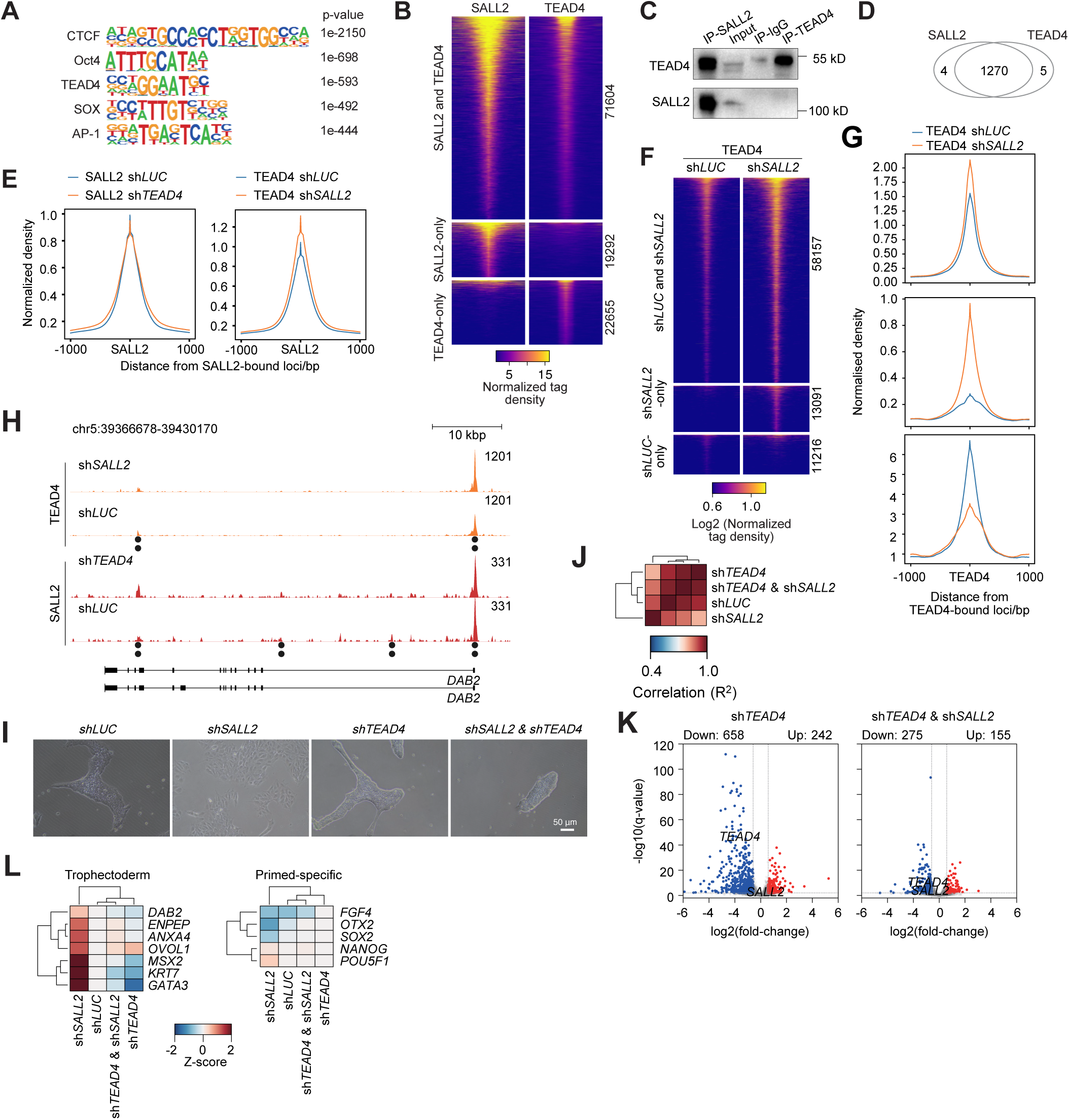
SALL2 interacts with and is bound to TEAD4-loci. **A** Transcription factor motif discovery at SALL2-bound loci. Selected motifs are shown. **B** Heatmap of CUT&Tag data for SALL2 and TEAD4 binding in hPSCs. TEAD4 ChIP-seq data is from GSE32465 ^52^, and GSE99202 ^53^. Peaks are centered on the overlap of SALL2 and TEAD4 binding loci, and the flanking 1 kbp. **C** Western blot of SALL2 or TEAD4 in a Co-IP of SALL2 and TEAD4 in hPSCs. **D** Venn diagram of SALL2 and TEAD4 Co-IP/MS identified proteins. **E** Pileup of CUT&Tag of SALL2 and TEAD4 in hPSCs transfected with an shRNA targeting *TEAD4*, *SALL2* or *LUC* as a control. Peaks are centered on SALL2 binding loci, and the flanking 1 kbp. **F** Heatmap of TEAD4 binding in hPSCs transfected with an shRNA targeting *LUC* as a control or an shRNA targeting *SALL2*. Peaks are centered on TEAD4 binding loci, and the flanking 1 kbp. **G** Pileup of CUT&Tag of TEAD4 binding in hPSCs transfected with an shRNA targeting *SALL2* or *LUC* as a control. Peaks are centered on TEAD4 binding loci, and the flanking 1 kbp. **H** Genome view of SALL2 and TEAD4 bound to *DAB2* TE-related locus in hPSCs transfected with an shRNA targeting *SALL2* or *TEAD4* or *LUC* as a control. **I** Morphology of hPSCs transfected with shRNAs targeting *SALL2*, *TEAD4*, both shRNAs or sh*LUC* as a control. **J** Cross-correlation heatmaps of the R^2^ correlation between all pair-wise samples. Samples are form hPSCs transfected with an shRNA targeting *TEAD4*, *SALL2*, both shRNAs or *LUC* as a control. **K** Volcano plots of the number of differentially regulated genes in the *TEAD4* hPSC knockdown, or in the dual sh*SALL2* and sh*TEAD4* knockdown cells. **L** Heatmaps of the expression of TE-related and pluripotent primed-specific genes in hPSCs transfected with an shRNA targeting *TEAD4*, *SALL2*, both shRNAs or *LUC* as a control.

As *TEAD4* is expressed in both hPSCs and hTSCs, whilst *SALL2* is high in hPSCs and low in hTSCs, we reasoned SALL2 may inhibit TEAD4 to impair TE differentiation. To understand the regulatory role of SALL2 we performed C&T for SALL2 and TEAD4 in control knockdowns (sh*LUC*) and when their opposite factor was knocked down (sh*SALL2* or sh*TEAD4*). Pileups at the SALL2 and TEAD4-bound loci showed a substantial overlap between the two factors (**Figure 3E-G and Supplementary Figure 5E**). Interestingly, the knockdown of *TEAD4* did not affect SALL2 binding, but *SALL2* knockdown led to increased TEAD4 binding (**Figure 3E**). This pattern could be exemplified by the pattern of binding at the TE-related *DAB2* and *TFAP2C* loci (**Figure 3H and Supplementary Figure 5F**). Finally, we confirmed the requirement of TEAD4 for the *SALL2* knockdown phenotype. Knockdown of *TEAD4* did not affect morphology and overall gene expression was similar to controls (**Figure 3I, J**) as previously reported for *TEAD4* knockout hPSCs ^29^. There were, however, a few hundred genes up and down-regulated, although GO analysis indicated the gene changes were not related to TE (**Figure 3K, L**). Instead, they appeared to bias the cells towards the immune system (**Supplementary Figure 5G**). In contrast to the *SALL2* knockdown, when both *SALL2* and *TEAD4* were knocked down the cells retained a normal hPSC morphology (**Figure 3I**). Their overall gene expression resembled controls (**Figure 3J**), they failed to up-regulate TE genes (**Figure 3L**) and did not downregulate pluripotency genes (**Figure 3L**). Indeed, the overall gene expression changes on sh*TEAD4*&sh*SALL2* knockdown resembled the control or sh*TEAD4* knockdown cells (**Figure 3J and Supplementary Figure 5H**). These data suggest TEAD4 is an essential requirement for the downstream effects of *SALL2* knockdown, and SALL2 is responsible for inhibiting TEAD4 activity in hPSCs to block TE gene expression.

### SALL2 maintains bivalent poised TE-related genes in primed cells

*TEAD4* is expressed at high levels in primed hPSCs but does not cause TE differentiation. As SALL2 is co-bound to the same loci as TEAD4, we reasoned that SALL2 must inhibit TEAD4 epigenetically. We first computationally analyzed the chromatin patterns at SALL2 bound loci using ChromHMM ^33^. ChromHMM predicted that SALL2-bound loci were a mixture of enhancers, TSSs and bivalent chromatin (**Figure 4A**). Indeed, reanalysis of hPSC epigenetic ChIP-seq data supported this assessment as SALL2-bound loci were rich in the active marks H3K4me3, H3K4me1, H3K27ac, but they also had appreciable levels of the inhibitory H3K27me3 (**Figure 4B**). To understand the epigenetic regulation, we analyzed our Co-IP/MS data for epigenetic regulators. SALL2 and TEAD4 co-precipitated a complex mixture of epigenetic complexes, including repressor and activator complexes (**Supplementary Table 1**). Particularly interesting was a mixture of activator and repressor complexes that included members of PRC2, SIN3A complexes, along with the BAF complex and activatory complexes, such as a HAT (ELP3) and COMPASS (**Figure 4C and Supplementary Figure 6A**). These interactions suggested the presence of several protein complexes; hence we reanalyzed hPSC epigenetic ChIP-seq data at SALL2 and TEAD4-bound loci. Interestingly, the repressive H3K27me3-catalysing PRC2 complex (exemplified by EZH2), the repressive histone-deacetylase SIN3A co-repressor complex (exemplified by SIN3A and HDAC2) and the activatory H3K4me3-related COMPASS (exemplified by RBBP5) were co-bound at SALL2 loci (**Figure 4D and Supplementary Figure 6B**). Overall, the interaction data suggests that SALL2 recruits a mixture of epigenetic regulatory complexes by binding directly to subunits to recruit the wider epigenetic regulatory complex to DNA and nucleosomes. This leads to the establishment of bivalent active/repressive chromatin at SALL2/TEAD4-bound loci.

**Figure 4.**
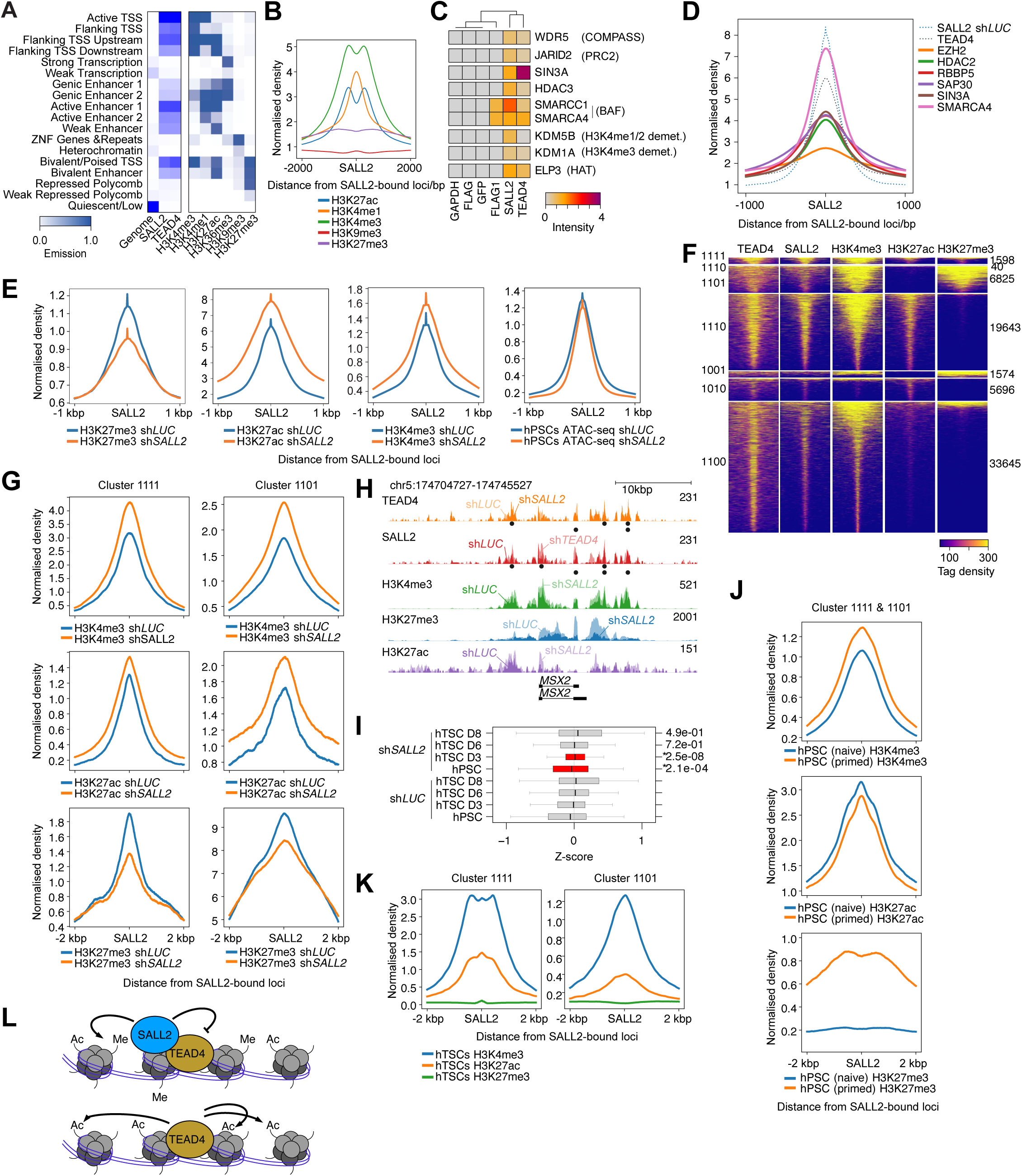
SALL2 regulates bivalent histone methylation/acetylation levels at TE-related genes. **A** ChromHMM of SALL2 binding. ChromHMM uses the 15-state model defined in ^54^. **B** Pileups of selected histone modifications at SALL2-bound loci in primed hPSCs. Data is from GSE59434 ^55^. **C** Heatmap of the mass spectrometry intensity for SALL2 or TEAD4 precipitated proteins for selected epigenetic factors. GAPDH, FLAG and GFP antibodies are shown as controls. **D** Pileups of selected epigenetic factors at SALL2-bound loci. Data is from GSE33281 ^56^, GSE29611 ^52^, GSE69647 ^57^, GSE22767 ^58^, and GSE24447 ^59^. **E** Pileups of H3K27me3, H3K27ac, H3K4me3 and ATAC-seq data at SALL2-bound loci in hPSCs transfected with shRNAs against *LUC* or *SALL2*. **F** Clustered heatmap of epigenetic states for H3K4me3, H3K27ac and H3K27me3 at all SALL2-bound loci in hPSCs. The binary cluster designations are indicated on the left and the number of peaks in each cluster on the right. **G** Epigenetic profiles at cluster 1111 (SALL2, H3K4me3, H3K27ac and H3K27me3 marked/bound loci) and cluster 1101 (SALL2, H3K4me3 and H3K27me3 bound/marked loci) in hPSCs transfected with shRNAs targeting SALL2 or LUC as a control. **H** Genome view of the TE-related gene MSX2 showing the changes in epigenetic profiles in hPSCs transfected with an shRNA targeting *SALL2* or *LUC*. **I** Boxplots of the expression of all genes with a TSS within 1000 bp of a SALL2 binding site in cluster 1111 and 1101. Expression is from hPSCs differentiated to hTSCs and transfected with an shRNA targeting *SALL2* or *LUC*. **J** Pileup of H3K27me3 in hPSCs and in hTSCs. hTSC data is from GSE193621^10^, and GSE266194 ^60^. **K** Epigenetic marks in primed and naive hPSCs at clusters 1111 and 1101. Naïve epigenetic ChIP-seq data is from GSE69647 ^57^,. **L** Schematic of the model of epigenetic regulation of TEAD4 by SALL2 at TE-related genes.

To explore changes in epigenetic regulation caused by SALL2 we performed ATAC-seq and CUT&Tag in hPSCs transfected with shRNAs targeting *SALL2* or *LUC* for the repressive epigenetic mark H3K27me3, and active mark H3K27ac and the promoter mark H3K4me3. In the *SALL2* knockdowns, ATAC-seq was modestly decreased, H3K27me3 decreased strongly, whilst H3K27ac and H3K4me3 increased at SALL2-bound loci (**Figure 4E**). This suggests knocking down *SALL2* reduced repressive and resolved bivalent chromatin to an active state. To explore in more detail, we divided the SALL2-bound loci into active (H3K27ac) and inhibitory (H3K27me3) clusters (**Figure 4F**). H3K4me3 marked the majority of loci, followed by the combination of H3K27me3 and H3K27ac (**Figure 4F**). Bivalent/repressive loci (including H3K27me3) numbered around 7000 loci. These loci also showed the most change on *SALL2* knockdown: reduced H3K27me3 and increased in H3K27ac and H3K4me3 (**Figure 4G**), indicating the chromatin is losing repression and gaining activation. The changes can be exemplified at specific TE genes, such as DAB2, GATA3, CDX2 and TBX3 (**Figure 4H and Supplementary Figure 6C**), and how they begin to adopt the epigenetic profile present in hTSCs when *SALL2* is knocked down (**Supplementary Figure 6D**). Importantly, the genes associated with Clusters 1111 (i.e. bivalent with H3K4me3, H3K27ac and H3K27me3) and 1101 (i.e. bivalent/repressive H3K4me3 and H3K27me3), showed increased gene expression in the early stages of TE differentiation (**Figure 4I**), suggesting that knocking down SALL2 enables TEAD4 activity mediated by histone modifications.

Naïve hPSC have low *SALL2* and this suggested that reduced *SALL2* in naïve cells preferentially benefits hTSC differentiation. This predicts that the TE bivalent loci would be weakened in naïve cells due to reduced SALL2 repression of TEAD4. Indeed, and epigenetic data for the bivalent loci highlights the complete loss of H3K27me3 in the naïve hPSCs (**Figure 4J**). Importantly, these epigenetic changes presage later epigenetic changes in hTSCs, as clusters 1111 and 1101 completely lack H3K27me3 in hTSCs (**Figure 4K**). Overall, this suggests that SALL2 inhibits TEAD4 activity in the primed state by establishing and maintaining bivalent chromatin to suppress TE fate (**Figure 4L**).

### Knockdown of *SALL2* improves TE commitment at the expense of epiblast in minimal media

We next sought to explore the role of SALL2 and TEAD4 in embryogenesis. Using a panel of early embryonic *in vitro* and *in vivo* stages showed how SALL2 is specific to hPSCs, and particularly primed, intermediate and E8-medium grown hPSCs (**Figure 5A**). SALL2 is otherwise high only in the ectoderm, which agrees with our experiments showing the overexpression of SALL2 promotes ectoderm (**Figure 1P**). *TEAD4*, conversely, is high during development, but reduced in the primed state (but is still detectable) and is then lost in somatic differentiation (**Figure 5A**). This is clearly illustrated in scRNA-seq from pre-implantation human embryos ^34^, which shows how *TEAD4* is high throughout the pre-implantation and extraembryonic tissues, is briefly co-expressed with *SALL2*, but is then lost from subsequent somatic cells (**Figure 5B**). This supports a role for SALL2 suppressing TEAD4 in the epiblast.

**Figure 5.**
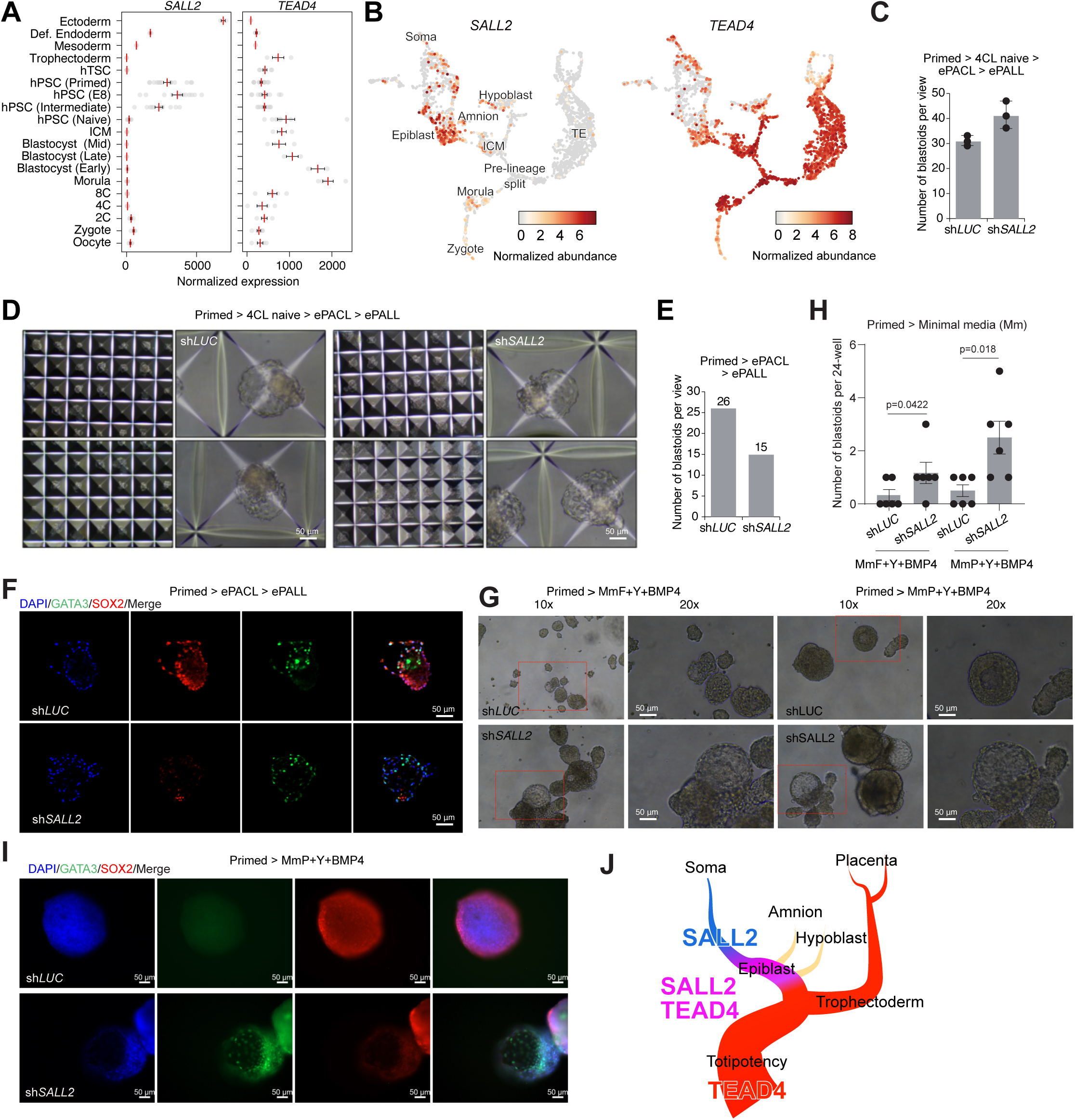
Reduced *SALL2* promotes blastoid formation, but at the expense of an epiblast. **A** Expression of SALL2 and TEAD4 in a panel of RNA-seq data from pre-implantation and early post-implantation stages. Data is from the reanalysis performed in Refs. ^43^ and other data from GSE93241 ^61^, GSE75868, GSE85689 ^62^, PRJNA383735 ^63^, SRP115256 ^64^, CNP0001454 ^65^, and GSE242351 ^18^. **B** UMAPs for the expression of *SALL2* (left) and *TEAD4* (right) in human embryo development. Selected lineages are indicated. Data is taken from ^34^. **C** Bar chart of the number of blastoids generated using the 4CL-naïve to ePACL then ePALL protocol. Experiment was performed three times. **D** Bright-field microscopy of blastoid transfected with a control shRNA (shLUC) or with an shRNA targeting *SALL2*. Scale bar, 100 µm. The experiment was repeated three times with similar results. **E** Bar chart of the number of blastoids generated using a Primed hPSCs to ePACL then ePALL based protocol. The experiment was performed once. **F** Immunostaining of GATA3 and SOX2 in blastoids generated using a primed hPSC to ePACL then ePALL blastoid protocol. Scale bars = 50 μm. **G** Morphology of blastoids generated in Minimal Media Five (MmF) based on 5iLAF medium, or Minimal Media P (MmP) based on PACL/PALL medium, with sh*SALL2* knockdown or LUC as a control. **H** Bar plots of the number of blastoids generated in Minimal Media Five (MmF) based on 5iLAF medium, or Minimal Media P (MmP) based on PACL/PALL medium, with sh*SALL2* knockdown or LUC as a control. **I** Representative IF images of 120 h blastoids transfected with sh*SALL2* or sh*LUC*. Blastoids were stained for GATA3 (green) and SOX2 (red). Shown is the maximum projection. Scale bars = 50 μm. **J** Model of the developmental interaction between SALL2 and TEAD4 in human embryonic development.

Ideally, we would next knockdown *SALL2* in human embryos with the prediction that at gastrulation they would have an absent or impaired epiblast. However, this experiment is difficult due to the ethical issues and paucity of embryos available for experimentation. Blastoids have emerged as a powerful model of early human embryonic development ^35–39^. Mechanistically blastoid derivations generally start with naïve hPSCs that transdifferentiate to TE. This suggests one limiting factor is the efficiency of transdifferentiation of TE cells. Indeed, blastoids tend to contain a larger number of epiblast-like cells, and a reduced number of TE cells, compared to a normal blastocyst ^34^. As *SALL2* KD improves TE differentiation, we reasoned it could potentially generate blastoids with improved TE. We first tried a protocol based on a brief conversion of primed hPSCs to naïve using 4CL-naive medium, followed by sequential culture in ePACL and ePALL medium ^40^. There was a modest increase in the number of blastoids when *SALL2* was knocked down (**Figure 5C**), however there was no visible change in the blastoid’s morphology (**Figure 5D**). Considering that the naïve state already has low *SALL2* (**Figure 5A**), this suggests that if blastoids are derived from naïve hPSCs they will naturally downregulate *SALL2* leading to no further effect when we concurrently knocked down *SALL2*. Consequently, we explored blastoids generated directly from primed hPSCs. Initially, we just omitted the 4CL-naive medium stage and cultured in ePACL then ePALL. Knockdown of *SALL2* under these conditions reduced the number of blastoids (**Figure 5E**), but increased TE content, as identified by GATA3 immunostaining (**Figure 5F**). However, morphological differences between the sh*LUC* and sh*SALL2* blastoids were overall modest (**Supplementary Figure 7A**). Potentially this is due to the ePACL medium containing PD0325901, CHIR99021, and LIF that may induce a transient naïve state.

Hence, we investigated blastoid generation using minimal media containing the smallest number of exogenous cytokines and inhibitors. We performed a small screen starting with the base medium used in 5iLAF blastoid generation (i.e. the 5iLAF medium with no cytokines or inhibitors; MmF), and we added back the inhibitors or growth factors until we observed the generation of cavitated blastoid-like structures (**Supplementary Figure 7B**). Under most conditions, whilst spheres formed, there was no signs of cavitation. The exception was cells grown in minimal media (MmF), or the minimal media from PACL/PALL (MmP) supplemented with the ROCK inhibitor Y-27632 and BMP4 (**Figure 5H**). Under these conditions, and with the knockdown of *SALL2*, we observed cavitated blastoid-like structures that were positive for GATA3 and had a small SOX2+ population of cells (**Figure 5I**). These data suggest that low *SALL2* can overcome barriers to blastoid generation in the absence of conversion to a naïve state. Based on this data, we propose a model whereby TE is the default fate of blastomeres, as TEAD4 is only downregulated when *SALL2* is co-expressed at the epiblast stage that causes cells to commit to the soma (**Figure 5L**).

## Discussion

Lineage decisions are complex processes of bifurcated decisions. Here, we show how SALL2 impairs TEAD4 activity to specify TE differentiation. Our data suggests SALL2 acts as an epigenetic lock on TEAD4 activity by blocking it from specifying TE and causing the reversion of ICM/epiblast cells to a TE-like fate. The timing of this inhibition is key to this idea. TEAD4 is expressed in all stages up to the epiblast, whilst SALL2 is low, and begins expression at the ICM but reaches a peak at the epiblast. Importantly, the point of dual-expression of SALL2 and TEAD4 is in the ICM and epiblast, which is after the lineage split to the ICM and TE. This suggests SALL2 and TEAD4 act to stop reversion or trans-differentiation to the default state of TE. Recently the Niakan research group published similar results in a preprint that also support our findings ^41^. Briefly, they found that knocking out *SALL2* also results in the emergence of a TE-like phenotype. They also identified a bias to primitive endoderm, which we did not observe, but this may be caused by their use of naïve *SALL2* knockout hPSCs which may retain primitive endoderm capability. Interestingly, they showed how SALL2 suppresses WNT, FGF2 and TGFB/BMP4 signaling to block TE differentiation. This may help explain why our blastoids can omit FGF2 and CHIR99021. However, why blastoids require complex media that includes a large and diverse number of inhibitors and exogenous signaling factors ^35,36,39^, whilst normal human embryos cultured in vitro require no exogenous inhibitors or cytokines ^42,43^, remains unclear. Potentially improved TE differentiation by modulating the pathways up and downstream of SALL2 may help improve blastoids.

Mouse Sall2^-/-^ embryos give rise to phenotypically normal animals ^44^, although in some backgrounds they show neural tube defects ^45^. This suggests differences between mouse and human TE specification. *Sall2* in mouse is similarly expressed at the epiblast stage ^45^, but potentially mouse TEAD4 performs different roles, as *Tead4*^-/-^ mice arrest at the morula stage. Indeed, the roles and timing of the key TE factors TEAD4, GATA3 and CDX2 are divergent in mouse and human ^29^, with CDX2 dominant in mouse ^8^, but GATA3 more important in human ^29^. Indeed, hTSCs have low *CDX2*, but high *GATA3* and *TEAD4* (and low *SALL2*) ^19^. Interestingly, bovine knockouts of *TEAD4* develop normally to the blastocyst stage ^46^, supporting divergent roles for these factors in different species.

Weakening histone acetylation is a feature required for TE differentiation, for example by modulating acetyl-CoA availability through metabolic changes leading to excess alpha-ketoglutarate ^47^. This weakened acetylation disrupts the pluripotency network, allowing the cells efficiently transdifferentiate to hTSCs. However, this does not explain how naïve hPSCs, which have surprisingly high levels of H3K27me3 ^48^, are more permissive to hTSC differentiation than primed hPSCs. Our data suggests SALL2 is a key distinguishing factor: low in naïve and high in primed hPSCs. Indeed, the high H3K27me3 and bivalency (maintained by SALL2) in hPSCs must be downregulated for differentiation to and from hTSCs ^49,50^.

Overall, our data suggests that in primed hPSCs SALL2 inhibits TEAD4 activity by establishing bivalent chromatin. Upon the loss of SALL2, either through SALL2 knockdown or during differentiation into hTSCs, TEAD4 becomes activated and bivalent inhibitory chromatin is resolved into an active chromatin state, thereby promoting TE differentiation. The biopotential switch of SALL2/TEAD4 is reminiscent of the OCT4/CDX2 switch that specifies the ICM and the TE in mouse ^8^, as well as the Tead4/Tfap2c switch ^51^. Our data suggests SALL2 functions as a later developmental safeguard within this regulatory framework, preventing reversion to the TE lineage after primed pluripotency has been established. Mechanistically, SALL2 epigenetically restrains TEAD4, promoting inhibitory chromatin and thereby reinforcing stable commitment to the somatic lineage. Overall, our data shows how SALL2 impairs TEAD4 activity to prevent trophoblast transdifferentiation and preserve the identity of primed hPSCs.

## Limitations of this study

We suggest that low *SALL2* is responsible for naïve hPSCs ability to differentiate to TE, but *TEAD4* is expressed in both naïve and primed hPSCs. Our model would thus predict that high TEAD4 in naïve hPSCs would cause the cells to spontaneously differentiate to TE as SALL2 no longer suppresses TEAD4. That naïve hPSCs do not differentiate to TE indicates other factors are inhibiting TEAD4 activity in naïve hPSCs. Similarly, TEAD4 cannot be solely responsible for cells defaulting to TE, as SALL2 knockdown causes only a small proportion of cells to differentiate to TE-like fate. Other factors must be present in the naïve state that promote the escape from a TE fate and other positive factors must push early epiblast cells to activate SALL2 to then inhibit TEAD4 for somatic lineage development. The identity of these factors is unknown. Finally, the blastoids generated here still require exogenous signaling, and are not efficiently generated, also confirming that SALL2 and TEAD4 are not the only key players in the bifurcation from blastomere to TE and epiblast.

## Acknowledgments

We thank Kathy Niakan and Huang Qiulin for discussions. We acknowledge the assistance of SUSTech Core Research Facilities. Funding was from the National Natural Science Foundation of China (W2512079 to A.P.H. 32370848 to M.A.E.), the National Key R&D Program of China (2025YFC3408900 to M.A.E.) the Guangdong Basic and Applied Basic Research Foundation (2023A1515111170 to XM.Z., 2025B1515020052 to DW.L), and the Science Technology and Innovation Commission of Shenzhen (RCBS20221008093109033 to XM.Z.).

## Author contributions

Q.Y. designed, performed and interpreted most of the experiments, prepared the figures and assisted in writing and revising the manuscript. XJ.F performed experiments, Z.L., HZ.Q. and ZT.L analyzed the data and revised the manuscript. XM.Z. revised the manuscript and acquired funding. G.M. GQ.T and M.A.E revised the manuscript and interpreted the data. A.P.H. designed the study, performed the bioinformatic analysis, acquired funding, supervised the study, and wrote the manuscript. All authors revised the manuscript.

## Data accessions

Data generated in this study is available under accession number GSE338016, GSE338017, GSE338018, GSE338019.

