## Supplementary Text and Figures for "SALL2 constrains TEAD4 by maintaining repressive chromatin to restrict trophectoderm identity"

[4]3DC STAR Lab, BGI CELL, Shenzhen, China

*Equal contribution.

#Correspondance:

**Supplementary Table 1**. Co-IP/MS identified proteins in SALL2 or TEAD4 immunoprecipitations from hPSCs.

#### Methods

##### hPSC culture, transfection, knockdowns

The human pluripotent stem cell (hPSC) line H1 (kindly provided by M. A. Esteban, Beijing Genomics Institute) and HEK293T cells (ATCC; RRID:CVCL_B488) were utilized in this study. hPSCs were maintained on plates pre-coated with Matrigel (354277, Corning) and cultured in mTeSR1 medium (85850, STEMCELL Technologies). The culture medium was replenished daily. For routine maintenance, hPSCs were passaged every 4–5 days via single-cell dissociation using Accutase (A6964, Sigma). HEK293T cells were cultured in DMEM-H basal medium supplemented with 10% fetal bovine serum (FBS, NATOCOR), with the medium exchanged every day.

For shRNA-mediated knockdown, the target sequences were cloned into the pLKO.1 puro vector (Addgene plasmid #8453). For SALL2 overexpression, the *SALL2* open reading frame (ORF) was subcloned into the pKD-Flag-IRES2-Puro and fused with the GFP-FLAG tag to express the SALL2-GFP-FLAG fusion protein. Lentiviral particles were generated by co-transfecting HEK293T cells with the respective target plasmid (knockdown or overexpression) along with the packaging plasmids pMD2.G and psPAX2 using polyethylenimine (PEI). Viral supernatants were harvested 48 hours post-transfection, purified, and either used immediately or stored at –80 °C for subsequent use. For lentiviral transduction, the purified viral particles were applied to human pluripotent stem cells (hPSCs). To ensure stable integration and establish semi-stable cell lines, the infected cells were subjected to puromycin selection beginning 48 hours after viral exposure.

#### hTSC differentiation

The differentiation of H1 cells into human trophoblast stem cells (hTSCs) was executed following a modified version of the protocol established by Okea *et al*[1]. Briefly, puromycin selected H1 cells were harvested and inoculated into Matrigel-pre-coated 6-well plates at a density of 1 × $10^5$ cells per well. The induction was performed in 2 mL of hTSC-specific induction medium, which consisted of DMEM/F12 basal medium (C11330500BT, Gibco) supplemented with the following components: 0.1mM β-mercaptoethanol (21985-023, Gibco), 0.2% FBS (FBS, NATOCOR), 0.5% Penicillin-Streptomycin (SV30010, HyClone), 0.3% BSA (B2064-100g, Sigma-Aldrich), 1% ITS-X supplement (C0345, Beyotime), 1.5 µg/ml L-ascorbic acid (Sigma, A8960), 50 ng/ml EGF (AF-100-15-100, PEPROTECH), 2 mM CHIR99021 (SML1046, Sigma-Aldrich), 0.5 Mm A83-01 (HY-10432, MedChemExpress), 1 mM SB431542 (301836-41-9, Selleck), 10 nM TSA (V900931, Vetec) and 5 µM Y27632 (HY-10583, MedChemExpress). Cultures were maintained in a humidified incubator at 37°C with 5% $CO_2$, and the induction medium was replenished on a daily basis. Upon reaching Day 6 of the induction period, the resulting cells were harvested or processed for subsequent experimental analyses.

#### Blastoid culture

Human primed embryonic stem cells (hESCs) were used to generate blastoids through either a naïve-mediated or a direct induction strategy.

For the naïve-mediated strategy, primed hESCs were dissociated into single cells using Accutase and seeded onto mitomycin C-treated mouse embryonic fibroblasts (MEFs). Cells were cultured in 4CL medium for 4 days under hypoxic conditions (5% $O_2$, 5% $CO_2$, 37°C) to induce the naïve pluripotent state. Naïve hESCs were subsequently dissociated into single cells with Accutase for 5 min at 37°C, harvested in DMEM/F-12, and centrifuged at 300 ×g for 3 min. The cell pellet was resuspended in fresh medium and pre-plated onto 0.2% gelatin-coated dishes for 30 min at 37°C to deplete residual MEFs. The non-adherent cells were collected, passed through a 40-µm cell strainer, counted, and transferred into AggreWell 400 (34415, STEMCELL TECHNOLOGIES) plates at a density of 1.5 × $10^4$ cells per well. Cells were cultured in ePACL medium for 3 days, followed by ePALL medium for an additional 6 days. Blastoids were typically observed from day 7 onward.

For the direct induction strategy, primed hESCs were dissociated into single cells using Accutase, harvested by centrifugation, resuspended in fresh medium, counted, and directly seeded into ultra-low attachment 6-well plates at a density of 1.5 × $10^5$ cells per well without naïve-state conversion. Cells were cultured in ePACL medium for 3 days followed by ePALL medium for 6 days under the same culture conditions, and blastoids were collected for subsequent analyses.

The 4CL medium Advanced DMEM/F 12 (12634028, Gibco) and Neurobasal (21103049, Gibco) mixed 1:1, supplemented with N2 (A1370701, Gibco), B27 (17504044, Gibco), non-essential amino acids (11140050, Gibco), GlutaMAX (A1286001, Gibco), sodium pyruvate (25-000-CI, Cellgro), penicillin/streptomycin (SV30010, HyClone), 10 nM DZNep (S7120, Selleck), 5 nM TSA (V900931, Vetec), 1 $\mu$M PD0325901 (1408, Axon), 5 $\mu$M IWR-1 (I0161, Sigma), 20 ng/ml human LIF (300-05, Peprotech), 20 ng/ml Activin A (120-14E, Peprotech), 50 $\mu$g/ml L-ascorbic acid (Sigma, A8960) and 0.2% (v/v) Matrigel.

The ePACL medium consisted of a 1:1 mixture of Advanced DMEM/F 12 (12634028, Gibco) and Neurobasal (21103049, Gibco) mixed 1:1, supplemented with N2 (A1370701, Gibco), B27 (17504044, Gibco), 1% GlutaMAX (A1286001, Gibco), 1% NEAA (11140050, Gibco), 1% sodium pyruvate (25-000-CI, Cellgro), 0.3 µM PD0325901 (1408, Axon), 100 ng/mL Activin A (120-14E, Peprotech), 10 ng/mL human LIF (300-05, Peprotech), 3 µM CHIR99021 (SML1046, Sigma-Aldrich), and the CEPT cocktail (HY-K1043, MedChemExpress).

The ePALL medium consisted of a 1:1 mixture of Advanced DMEM/F 12 (12634028, Gibco) and Neurobasal (21103049, Gibco) mixed 1:1, supplemented with N2 (A1370701, Gibco), B27 (17504044, Gibco), 1% GlutaMAX (A1286001, Gibco), 1% NEAA (11140050, Gibco), 1% sodium pyruvate (25-000-CI, Cellgro), 1 µM PD0325901 (1408, Axon), 1 µM A83-01 (HY-10432, MedChemExpress), 10 ng/mL human LIF (300-05, Peprotech), 0.5–1 μM lysophosphatidic acid (LPA) (HY-107614, MedChemExpress), and the CEPT cocktail (HY-K1043, MedChemExpress).

**Generation of human blastoid in modified minimal media**

Primed human pluripotent stem cells (hPSCs/hESCs) were dissociated into single cells by incubation with Accutase (Merck Sigma-Aldrich, A6964) for 3 minutes. Following centrifugation, the cells were harvested, resuspended in fresh minimal medium, and counted. Without undergoing naïve-state conversion, the single cells were directly seeded into ultra-low attachment 24-well plates at a density of $1.5 \times 10^4$ cells per well. These cells were then cultured in either MmF or MmP minimal media with 5 μM Y27632 (HY-10583, MedChemExpress), and 10 ng/ml BMP4 (ab87063, abcam) for 8 days, with the culture medium replenished every other day. Following this period, the resulting blastoids were collected for subsequent downstream analyses.

The MmF medium consisted of Advanced DMEM/F 12 (12634028, Gibco) and Neurobasal (21103049, Gibco) mixed 1:1, supplemented with N2 (A1370701, Gibco), B27 (17504044, Gibco), 0.5% KnockOut SR (10828028, Gibco), 1% GlutaMAX (A1286001, Gibco), 1% MEM Non-Essential Amino Acids (NEAA) (11140050, Gibco), 1% penicillin-streptomycin (SV30010, HyClone), 0.1 mM β-mercaptoethanol (21985-023, Gibco), 50 μg/ml BSA (B2064-100g, Sigma-Aldrich). Alternatively, the MmP medium comprised a 1:1 mixture of Advanced DMEM/F 12 (12634028, Gibco) and Neurobasal (21103049, Gibco) mixed 1:1, supplemented with N2 (A1370701, Gibco), B27 (17504044, Gibco), 1% GlutaMAX (A1286001, Gibco), 1% NEAA (11140050, Gibco), 1% sodium pyruvate (25-000-CI, Cellgro).

**Western Blot**

For Western blot analysis, total cellular proteins were harvested utilizing RIPA lysis buffer (P0013B, Beyotime), with their concentrations subsequently determined using a BCA protein assay kit (Thermo Fisher Scientific). Antibodies used include TUBG (1:1,000 dilution, 5886S, CST), SALL2 (1:1,000 dilution, A303-208A, Thermo Fisher Scientific) and TEAD4 (1:1,000 dilution, 12418-1-AP, Proteintech). The membranes were then incubated with an HRP-conjugated secondary antibody (1:2,000 dilution,

20818, Biotium), and the target protein bands were developed using the enhanced BeyoECL method (P0018AS, Beyotime) before being captured and visualized on a Tanon 6100C chemiluminescence imaging system.

**Immunofluorescence**

For immunofluorescence of adherent cells, cells were briefly rinsed with PBS (C10010500BT, Gibco) and fixed in 4% (w/v) paraformaldehyde (PFA) (P0099, Beyotime) for 5 min at room temperature (RT). Then cells were washed with PBS and permeabilized with 0.25% (v/v) Triton X-100 (T8787, Sigma) for 3 min at room temperature. The samples were then incubated with primary antibodies diluted in blocking buffer at 4 °C overnight. Subsequently, the cells were washed with a buffer consisting of 0.1% (v/v) Tween-20 in PBS (C10010500BT, Gibco) and treated with the corresponding Alexa Fluor secondary antibodies (5 µg ml–1, A-11034, A-11004 and A-21235, Life Technologies) in wash buffer for 2 h at room temperature. Nuclei were stained using 4,6-diamidino-2-phenylindole (DAPI) (5 µg ml–1, 62248, Thermo Fisher Scientific). Fluorescence signals were ultimately captured and visualized using a Zeiss LSM 980 confocal microscope.

For immunofluorescence of blastoids, samples were fixed with 4% (w/v) paraformaldehyde (PFA) (P0099, Beyotime) for 30 min at room temperature or overnight at 4 °C, followed by rinsing with Dulbecco's phosphate-buffered saline (DPBS, BasalMedia, B210KJ). Blastoids were permeabilized with 0.75% Triton X-100 (T8787, Sigma) in DPBS for 1 h using a mouth pipette, then incubated in blocking buffer (DPBS containing 3% BSA and 0.1% Triton X-100) for 1 h at RT. Blastoids were incubated with primary antibodies diluted in blocking buffer at 4 °C overnight. Wash for three times with DPBS containing 0.1% Triton X- 100. Subsequently, blastoids were incubated for 1 h at RT in the dark with Alexa Fluor (488, 555, or 647) conjugated secondary antibodies (Invitrogen) at a 1:1000 dilution. After secondary antibody incubation, cell nuclei were stained with 4',6-diamidino-2- phenylindole (DAPI, Sigma, D9542). Confocal images were acquired using Zeiss LSM800 or LSM880 laser scanning microscopes, and subsequent image processing, including diameter measurements, was performed using ZEN software.

**Co-immunoprecipitation and mass spectrometry**

Co-IP was performed using Immunoprecipitation Kit with Protein G Magnetic Beads (P2177S, Beyotime) according to the manufacturer's instructions. Briefly, cell lysates were prepared using the provided Lysis Buffer supplemented with the Protease Inhibitor Cocktail. Pre-washed Protein G magnetic beads were first incubated with the specific primary antibody or normal IgG control to form bead-antibody complexes. The prepared cell lysates were then added to these complexes and incubated overnight at 4°C with gentle rotation. Following magnetic separation, the beads were washed three times with Lysis Buffer to remove non-specifically bound proteins. Proteins were detected on an Agilent 7700X.

Analysis of Co-IP–MS was performed using MaxQuant (v.2.6.5.0), and processed using glbase3 [2]. In brief, peptides detected by MaxQuant were filtered based on several criteria. A protein was considered detected if it had at least one unique peptide and a minimum intensity of 1,000,000 (Razor + Unique). A protein was considered specific to SALL2 or TEAD4 if the intensity was at least two-fold above the anti-FLAG, GAPDH or GFP control. The resulting peptides were filtered to remove a list of common contaminating proteins (protein names starting with RPL, RPS, TUB, GAPDH, ACT, MRPS, sm-, MYH, FLN, MYO, APOA1, MYL, IGHG, IGLV, IGHV, IGKV, COL, KRT, EIF and ATP). Epigenetic factors or transcription factors were determined based on the EpiFactors and AnimalTFDB databases [3,4]. The resulting table of filtered proteins is in **Supplementary Table 1**.

**RNA extraction, RT-qPCR, bulk RNA-seq analysis**

Total RNA from cells was extracted using RNAzol RT (MRC, RN190) according to the manufacturer's protocols. cDNA synthesis by using a PrimeScript RT Master Mix (Takara, RR036A). Real-time PCR was performed in triplicate using SYBR Premix Ex Taq (Takara, RR820A) and using a Biorad Real-time PCR system. The primers used are listed in STAR Methods.

Bulk RNA-seq was analysed as described in [5]. Briefly, reads were aligned to the hg38 genome using STAR aligner [6] with the settings: --readFilesCommand zcat --outFilterMultimapNmax 100 --winAnchorMultimapNmax 100 --outMultimapperOrder Random --runRNGseed 777 --outSAMmultNmax 1 --outSAMtype BAM Unsorted --

twopassMode Basic --outFilterType BySJout --alignSJoverhangMin 8 --alignSJDBoverhangMin 1 --outFilterMismatchNmax 999 --alignIntronMin 20 --alignIntronMax 1000000 --alignMatesGapMax 1000000, and assigned to genes and TEs using te_counter (a reimplementation of scTE [7]; https://github.com/oaxiom/te_counter) with the settings: -m genes_tes and using the GENCODE v42 transcriptome assembly [8]. Data was normalized using EDASeq [9]. Differential gene expression was called using DESeq2 [10], and a gene was considered differentially expressed if it had a Bonferroni-Hochberg corrected p-value less than 0.01. Fold-change was also used to discriminate differentially regulated genes, and this varied depending upon the experiment and the expected magnitude of change. In all cases a minimum fold-change of 1.5 was used. Gene ontology analysis was performed using goseq [11], and GSEA with fgsea [12]. Other analysis was performed using glbase3 [2].

**Single-cell RNA-seq and analysis**

H1 hPSCs cells expressing shRNA targeting *SALL2* or control *LUC* shRNA were harvested and dissociated into single-cell suspensions. The suspensions were adjusted to a concentration of 1000 cells/ul with a viability of >80%. The cells from each group were then separately loaded onto the 10x Genomics Chromium Controller to generate single-cell gel beads-in-emulsion (GEMs). Single-cell RNA-seq libraries were constructed using the Chromium Single Cell 3′ Reagent Kits v3.1 (10x Genomics, Pleasanton, CA, USA) according to the manufacturer's instructions. Briefly, within the GEMs, cells were lysed and mRNA was reverse-transcribed into barcoded cDNA. The GEMs were then broken, and the pooled cDNA was amplified via PCR. Following enzymatic fragmentation and size selection, Illumina-compatible sequencing adapters and sample-specific indices were added. The quality and size distribution of the final libraries were assessed using Agilent 5400 (Agilent Technologies). Finally, the libraries were sequenced on an Illumina NovaSeq Xplus platform, targeting a depth of approximately 35,000 read pairs per cell, with a sequencing configuration of 28 bp for Read 1, 90 bp for Read 2, and 10 bp for the i7 index.

Single cell RNA-seq was analyzed as previously described [13]. Briefly, reads were aligned to the hg38 genome using STAR-solo [6] with the settings: --soloType

Droplet --soloFeatures Gene Velocyto --soloBarcodeReadLength 0 --outFilterMultimapNmax 100 --winAnchorMultimapNmax 100 --outSAMmultNmax 1 --outSAMtype BAM Unsorted --twopassMode Basic --runRNGseed 42 --runThreadN 16 --readFilesCommand zcat --limitSjdbInsertNsj 4000000 --limitOutSJcollapsed 5000000 --soloUMIlen 12 --soloCBlen 16 --soloUMIstart 17, using an index generated from GENCODE v42 [8]. Reads were assigned to features using te_counter (a low memory compatible reimplementation of scTE) with the settings: --sc --strand [7]. Downstream analysis was performed using scanpy [14], batch correction by harmonypy [15], and assisted by sc_utils (https://github.com/oaxiom/sc_utils), and glbase3 [2].

#### CUT&Tag and ATAC-seq analysis

CUT&Tag was performed using the Hyperactive Universal CUT&Tag Assay Kit for Illumina Pro (TD904, Vazyme) according to the manufacturer's instructions. Briefly, using the kit, around 100,000 cells for each sample were collected and incubated with the Nuclear Extraction Buffer to extract nuclei, which were then incubated with the provided pre-activated ConA Beads Pro. Digitonin was used to permeate the cell membrane. Next, a primary antibody was diluted at 1:50 and incubated with the sample at 4℃ overnight. After centrifugation, the reaction solution was collected, incubated at room temperature with the corresponding anti-Rabbit secondary antibody (ABIN101961, Antibodies-Online) diluted 100 times, and put on a rotator for 1 h. After the supernatant was discarded and the pellet was cleaned with Dig-wash buffer, the nuclei-magnetic bead complex was generated and was then mixed with Hyperactive pA/G-Transposon Pro of which the final concentration was 40 nM at room temperature for 1 h. Trueprep Tagment Buffer L was added, transposons fused with Protein A/G, and targeted DNA sequences to add the splitter sequence to both ends of the cut fragment. After the sample was processed by pre-activated DNA Extract Beads Pro (TD904, Vazyme), PCR amplification with total cycle numbers of 10 was performed with the primers from DNA Adapter from TruePrep Index Kit V3 for Illumina (Vazyme #TD203). The amplified product was then cleaned using the VAHTS DNA Clean Beads, according to the manufacturer's instructions (Vazyme #N411). The resulting sample was sent for quality control and high-throughput sequencing on an Illumina sequencer.

ATAC-seq [16], was performed using the Hyperactive ATAC-Seq Library Prep Kit for Illumina (TD711, Vazyme), according to the manufacturer's instructions. Briefly, 100,000 cells were collected, and nuclei were extracted using pre-cooled Lysis Buffer. Next, according to the instructions, the cell genome was segmented by incubating at 37℃ for 30 min with the Tn5 complex, which carries a known DNA sequence tag. Then the DNA sequences with connectors were extracted using ATAC DNA Extract Beads (TD711, Vazyme). Afterward, PCR amplification with total cycle numbers of 11 was performed with the DNA Adapter primers from TruePrep Index Kit V3 for Illumina (Vazyme #TD203) to form a library. ATAC DNA Clean Beads (TD711, Vazyme) were used for two-step fragment sorting and product amplification, and finally, the samples were sent for library quality control and sequencing.

CUT&Tag and ATAC-seq data was analyzed as previously described [5,17]. Briefly, reads had adaptors trimmed aligned to the hg38 genome using bowtie2 [18] with the settings: -I 10 -X 1000 or -I 10 -X 2000 for ATAC-seq. Peaks were called using MACS2 [19] with a q-value of -q 0.01, and peak information shared using redefine_peaks [13]. Motif discovery was performed using HOMER [20]. Chromatin states were defined using ChromHMM [21] using the 15-state hPSC model defined in [22]. Other analysis was performed using glbase3 [2].

#### Lineage bias analysis

The lineage bias analysis was performed as described online and in a forthcoming publication (https://github.com/oaxiom/glbase_bio_data/tree/master/GSEA/Hs/lineage_bias). Briefly, RNA-seq data from the reanalysis in Refs. [23,24], and other data from GSE93241 [25], GSE75868, GSE85689 [26], PRJNA383735 [27], SRP115256 [28], CNP0001454 [29], GSE242351 [5] and GSE269273 [23] were used to generate panels of lineage/cell type-specific genes. Consequently, in the lineage specific measures a significant result for 'Primed' has no real meaning. Lineage-specific genes were determined as those significantly different (using DESEq2) versus the primed state. These sets of genes serve as a panel for measuring the differentiation bias of hPSCs. Significantly-associated lineages were determined using fgsea [12]. Only a positive and significant score has meaning in this analysis. The sets of lineage-specific genes are available online at: https://github.com/oaxiom/glbase_bio_data/tree/master/GSEA/Hs/lineage_bias/lineage_bias.gmt.

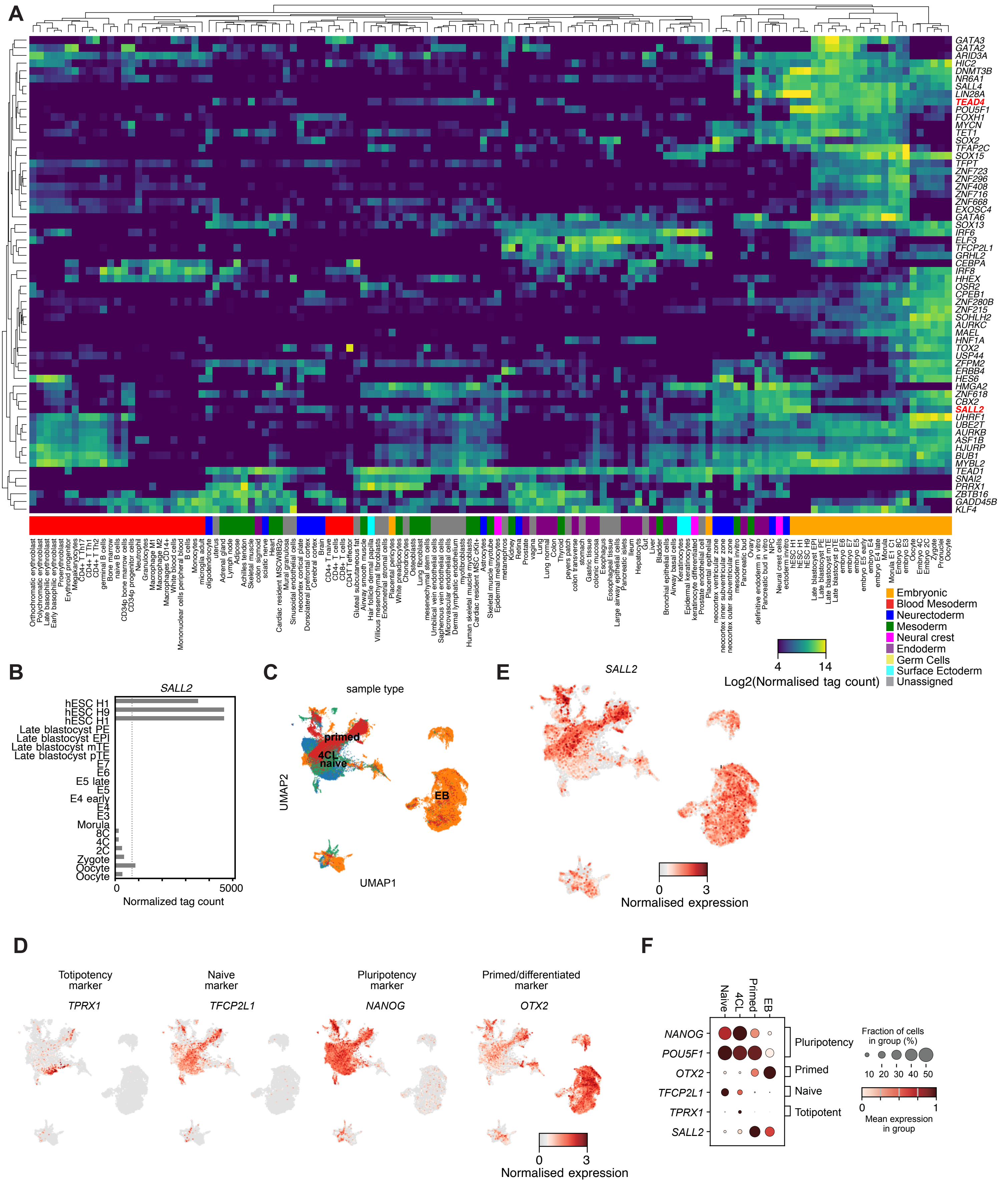


### Supplementary Figure 1

**Supplementary Figure 1. *SALL2* gene expression in early embryo and hPSC states**

- **A** Heatmap of selected transcription and epigenetic factors expressed in early embryonic development (orange). Data is from the reanalysis in Ref. [30].
- **B** Bar chart of the expression of *SALL2* gene during embryonic development and in primed hPSCs (H1, H9 lines). Data is from the reanalysis in Ref. [24].
- **C** UMAP of selected scRNA-seq data from primed, naïve (including 4CL cells), and EBs. Data is from GSE178274 [31], GSE166422 [32], GSE150311 [33], PRJNA631808 [30], and CNP0001454 [29].
- **D** UMAPs colored by the expression levels of the totipotent-related gene *TPRX1*, the naïve-specific gene *TFCP2L1*, the pluripotency marker *NANOG* and the primed/differentiated marker gene *OTX2*.
- **E** UMAP, as in panel C, colored by the expression of *SALL2*.
- **F** Bubble plot of selected pluripotency, naïve and primed genes and *SALL2*.

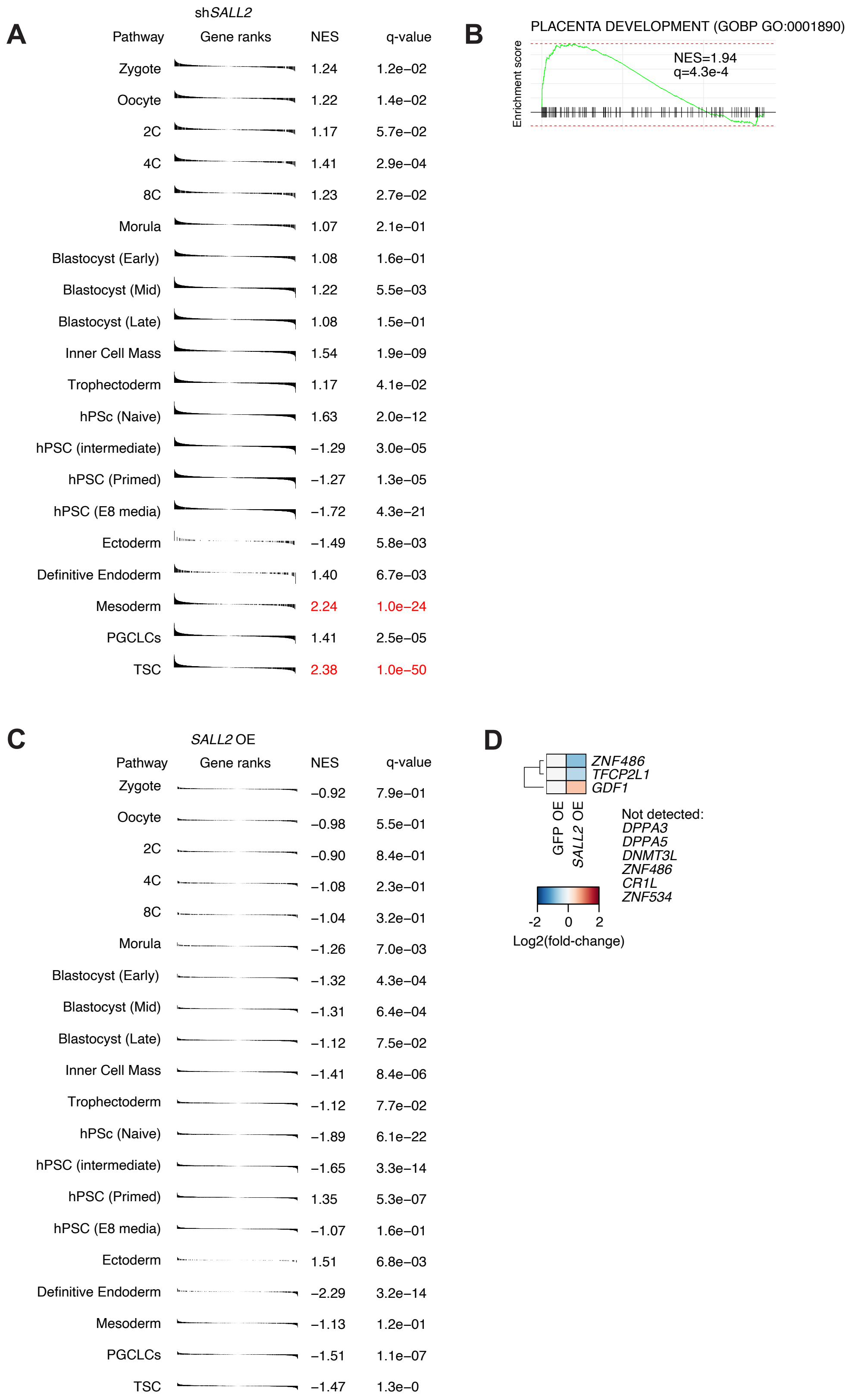


#### Supplementary Figure 2

**Supplementary Figure 2. Reduced *SALL2* promotes a TE-like cell fate.**

**A** GSEA maps for a panel of lineage-specific genes for the hPSCs transfected with an shRNA targeting SALL2 versus the control shLUC. The genes for each lineage are defined in https://github.com/oaxiom/glbase_bio_data/tree/master/GSEA/Hs/lineage_bias/lineage_bias.gmt. Up-regulated genes are on the left. NES = normalized enrichment score. q-value is the Bonferroni-Hochberg corrected p-value from fgsea.

**B** GSEA for GO BP category 'placenta development'.

**C** GSEA maps for a panel of lineage-specific genes for the hPSCs transfected with an vector overexpressing SALL2 versus a control vector containing GFP. The genes for each lineage are defined in https://github.com/oaxiom/glbase_bio_data/tree/master/GSEA/Hs/lineage_bias/lineage_bias.gmt. Up-regulated genes are on the left. Details are as in **panel A**.

**D** Heatmap of selected naïve-specific genes in the GFP or SALL2 overexpressing (OE) hPSCs. Note that DPPA3, DPPA5, DNMT3L, ZNF486, CR1L and ZNF534 were below the detection threshold.

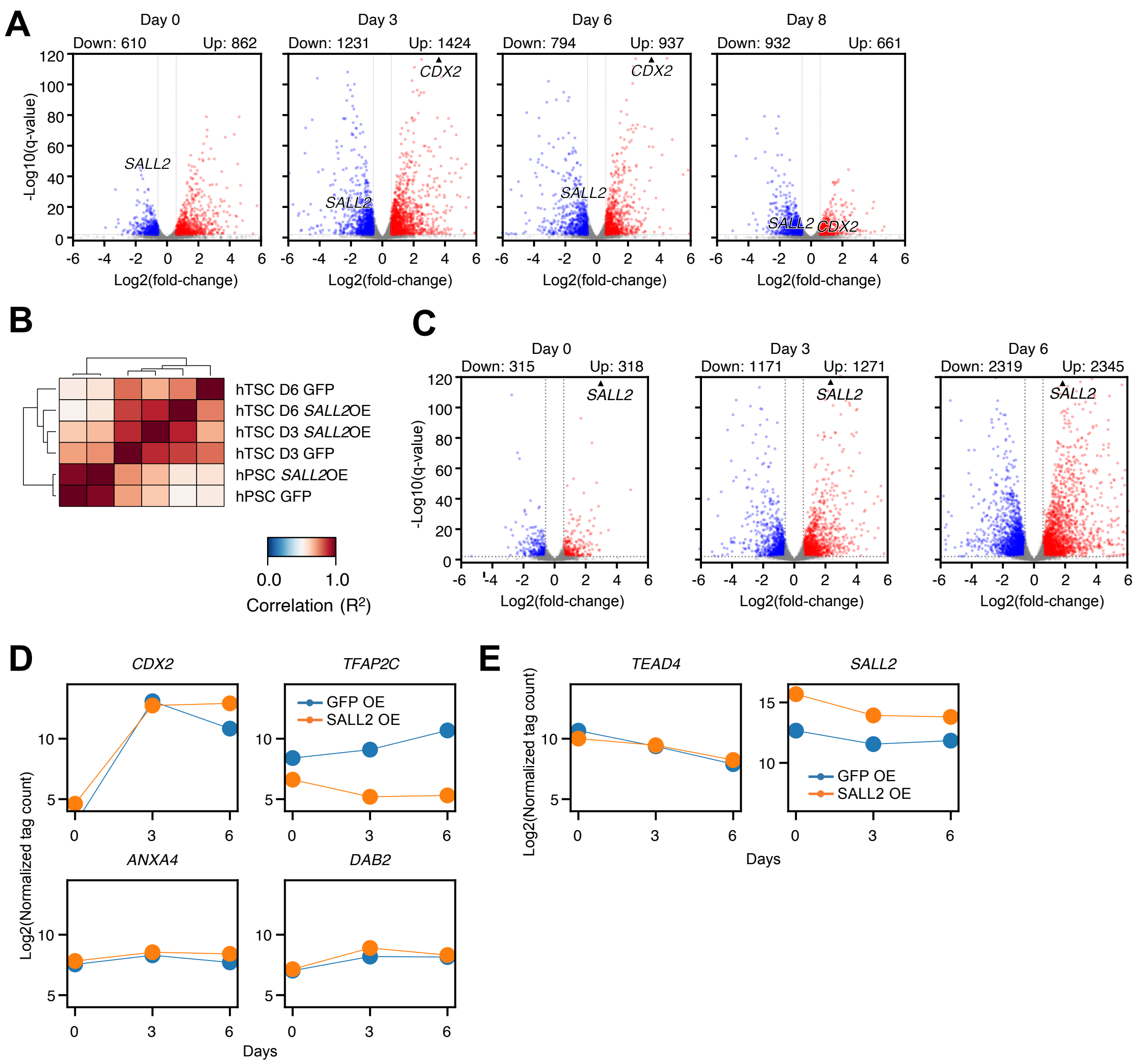


**Supplementary Figure 3**

**Supplementary Figure 3. Modulating *SALL2* improves hTSC differentiation.**

- **A** Volcano plot of day 0, 3, 6, 8 RNA-seq of the hTSC time course of cells transfected with shRNAs targeting shLUC or shSALL2.
- **B** Cross-correlation heatmap of hPSCs transfected with shRNAs against SALL2 or a control LUC.
- **C** Volcano plots of significantly differentially regulated genes at day 0, 3 and 6 in hPSCs differentiated to hTSCs.
- **D** Line plots showing the expression of selected trophectoderm genes in hPSCs overexpression *SALL2* or *GFP* (control).
- **E** Line plot of *SALL2* and *TEAD4* in hPSCs overexpressing *SALL2* or *GFP* (Control).

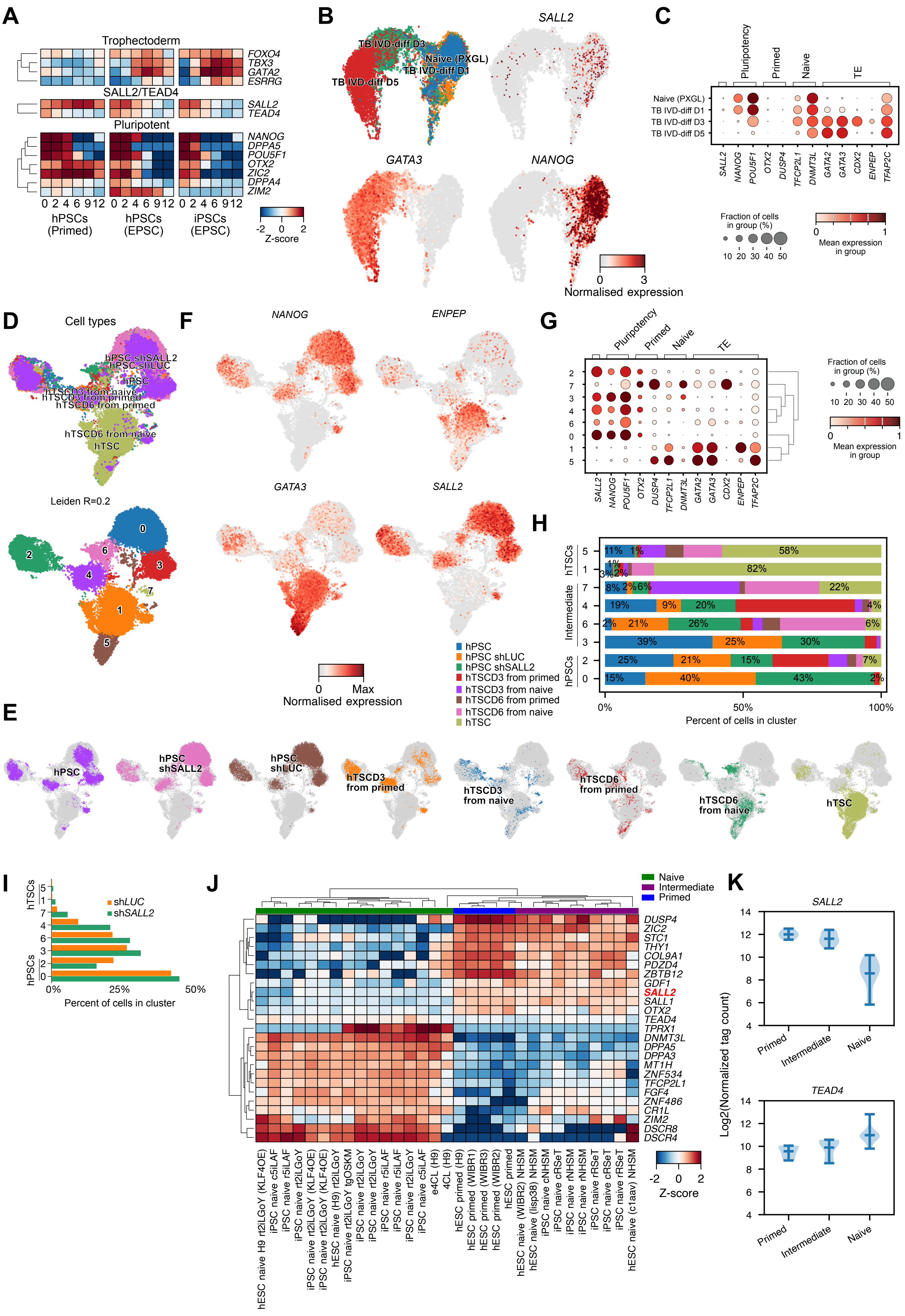


### Supplementary Figure 4

**Supplementary Figure 4. Naïve cells have reduced *SALL2* and downregulate it more efficiently in TE differentiation.**

**A** Heatmap of the expression of selected TE and pluripotency genes, and *SALL2* in primed hPSCs and in EPSCs derived from hPSCs or induced pluripotent stem cells (iPSCs). Data is from E-MTAB-7253 [34].

**B** scRNA-seq of a differentiation time course from PGXL naïve cells to TSCs. Data is from GSE166422 [32].

**C** Bubble plot of a selection of pluripotency, primed and naive-specific genes along with trophectoderm genes.

**D** UMAP plot for naïve and primed hPSCs differentiated to hTSCs (days 3 and 6), and embryo-derived hTSCs, along with hPSCs transfected with shRNAs targeting *SALL2* or *LUC* as a control. Data is from GSE281595 [35]. The upper panel is labelled by the sample identity and lower panel is clustered using the Leiden algorithm (resolution 0.2).

**E** UMAPs colored by the sample of origin.

**F** UMAPs of the hPSC primed and naïve differentiation to hTSCs colored by the expression of the *SALL2*, the pluripotency gene *NANOG*, and the TE-related genes *GATA3* and *EPAS1*.

**G** Bubble plot for a selection of hPSC primed, naïve-specific genes, trophectoderm-related genes and *SALL2*.

**H** Proportional bar chart showing the percentage of cells from each sample in the clusters defined in **panel D**.

**I** Bar chart of the percentage of hPSCs transfected with shRNAs targeting SALL2, or LUC as a control versus the scRNA-seq defined clusters from **panel D**.

**J** Heatmap of the expression of a selection of naïve and primed-specific genes and SALL2. Data is from the reanalysis performed in [5], based on GSE93241 [25], GSE75868, GSE85689 [26], PRJNA383735 [27], PRJNA397941 [28], and CNP0001454 [29].

**K** Violin plot of the expression of *SALL2* and *TEAD4* in the primed, intermediate and naïve cell types as defined in **panel I**.

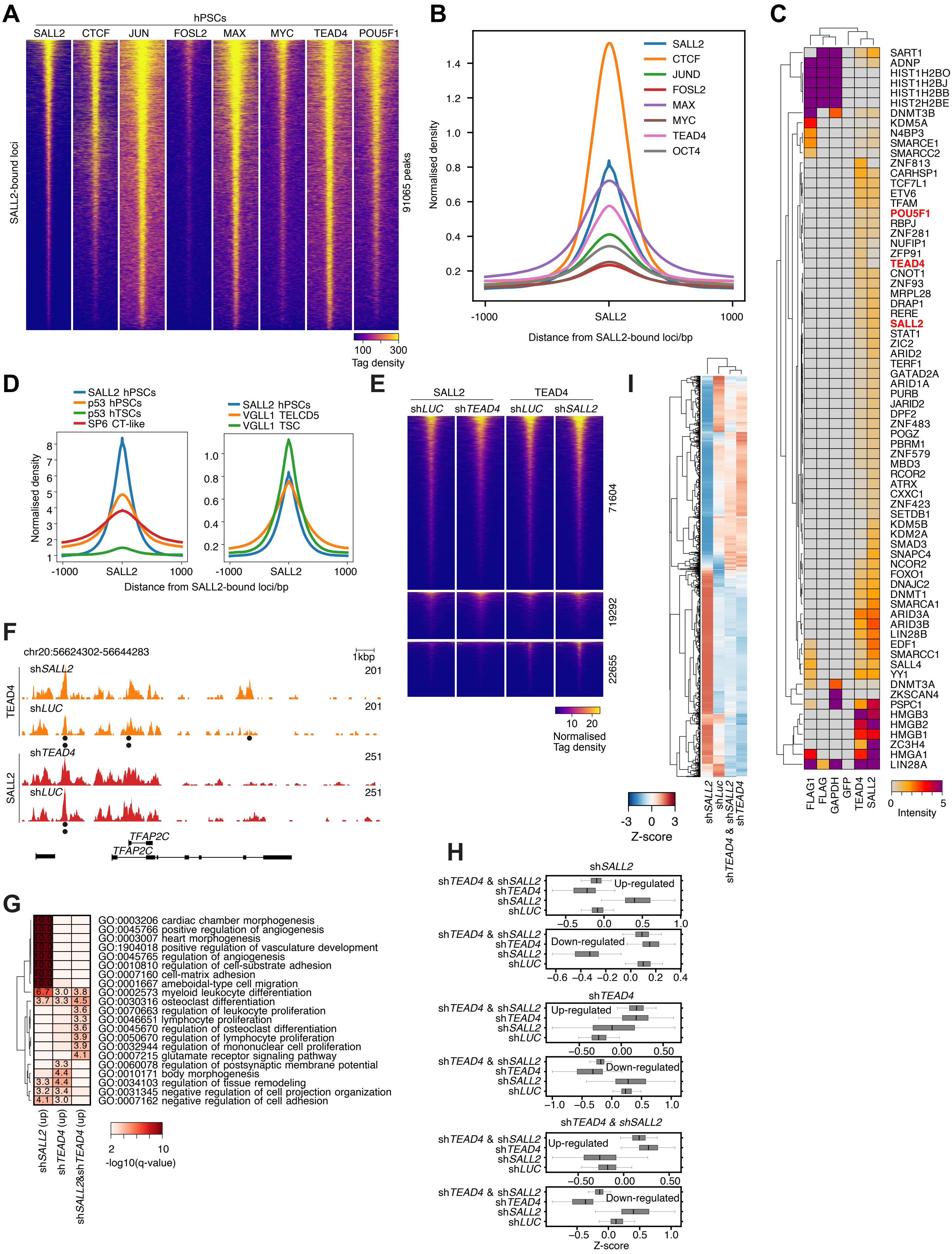


**Supplementary Figure 5**

**Supplementary Figure 5. SALL2 interacts with TEAD4.**

**A** Heatmap of selected transcription factors at SALL2-bound loci in hPSCs. Data is from GSE61475 [36], GSE52457 [37], GSE32465 [38], and this study (SALL2). Genes were selected based on their presence in the AnimalTFDB 3.0 database [4].

**B** Pileups of the heatmaps in **panel A**.

**C** Heatmap of the mass spec intensity score (/1e6) for TFs and epigenetic factors detected by mass spectrometry precipitated with SALL2 or TEAD4 or both. Controls (FLAG, FLAG1, GAPDH and GFP) are shown for comparison.

**D** Pileup of SALL2, and p53 in hPSCs and hTSCs, and SP6 in cytotrophoblast-like cells (left panel) and VGLL1 at SALL2-bound loci in hPSCs or hTSCs/TELCs (D5) (right panel). Data is from GSE277759 [39], GSE39912 [40], GSE236794 [41], and GSE193621[42].

**E** Heatmap of SALL2 and TEAD4 genome-wide binding in hPSCs transfected with shRNAs targeting LUC (control) or *SALL2* or *TEAD4*.

**F** Genome view of SALL2 and TEAD4 C&T data at the TE-related gene *TFAP2C*.

**G** Gene ontology of the differentially regulated genes in hPSCs transfected with shRNAs targeting SALL2, TEAD4, both or LUC as a control.

**H** Boxplots of the expression level of up and down-regulated genes in hPSCs transfected with shRNAs targeting *SALL2*, *TEAD4* or *LUC* as a Control.

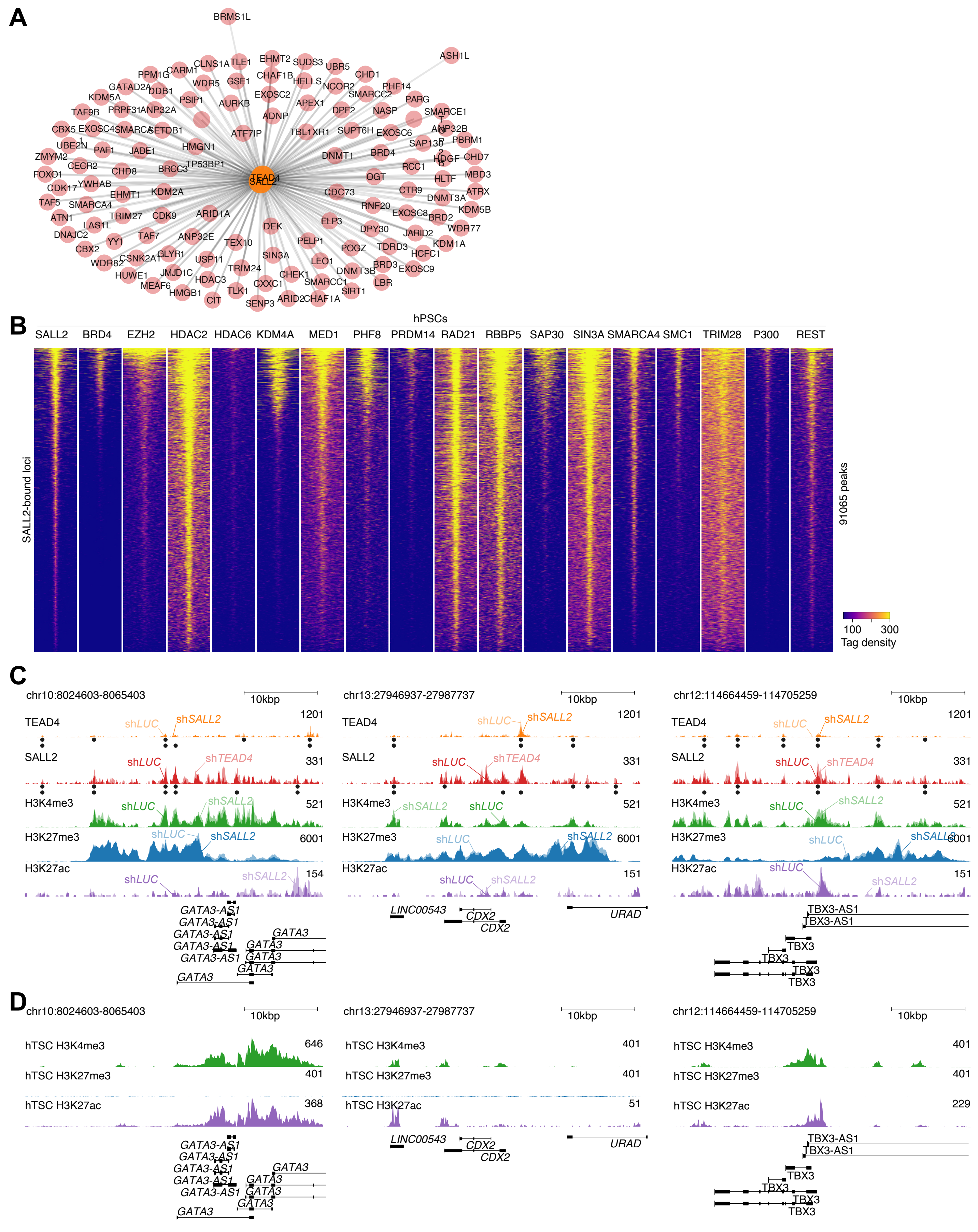


#### Supplementary Figure 6

**Supplementary Figure 6. Epigenetic co-factors for SALL2/TEAD4 in hPSCs.**

- **A** Network of nuclear-localized (defined as a score >=4 in the COMPARTMENTS database [43]) epigenetic factors co-bound with TEAD4 and SALL2 in the Co-IP/MS data. Epigenetic factors were defined based on their presence in the EpiFactors database [3].
- **B** Heatmaps of pileups at SALL2-bound loci in hPSCs for selected epigenetic factors. Data is from GSE33281 [44], GSE29611 [45], GSE69647 [46], GSE22767 [47], and GSE24447 [48].
- **C** Genome views for SALL2, TEAD4, H3K4me3, H3K27me3, and H3K27ac for selected TE-related genes in hPSCs transfected with shRNAs targeting *SALL2*, *TEAD4* or LUC as a control.
- **D** The same genome views as in **panel C**, but for the epigenetic histone modifications H3K4me3, H3K27me3 and H3K27ac in hTSCs. Data is from GSE193621[42], and GSE266194 [49].

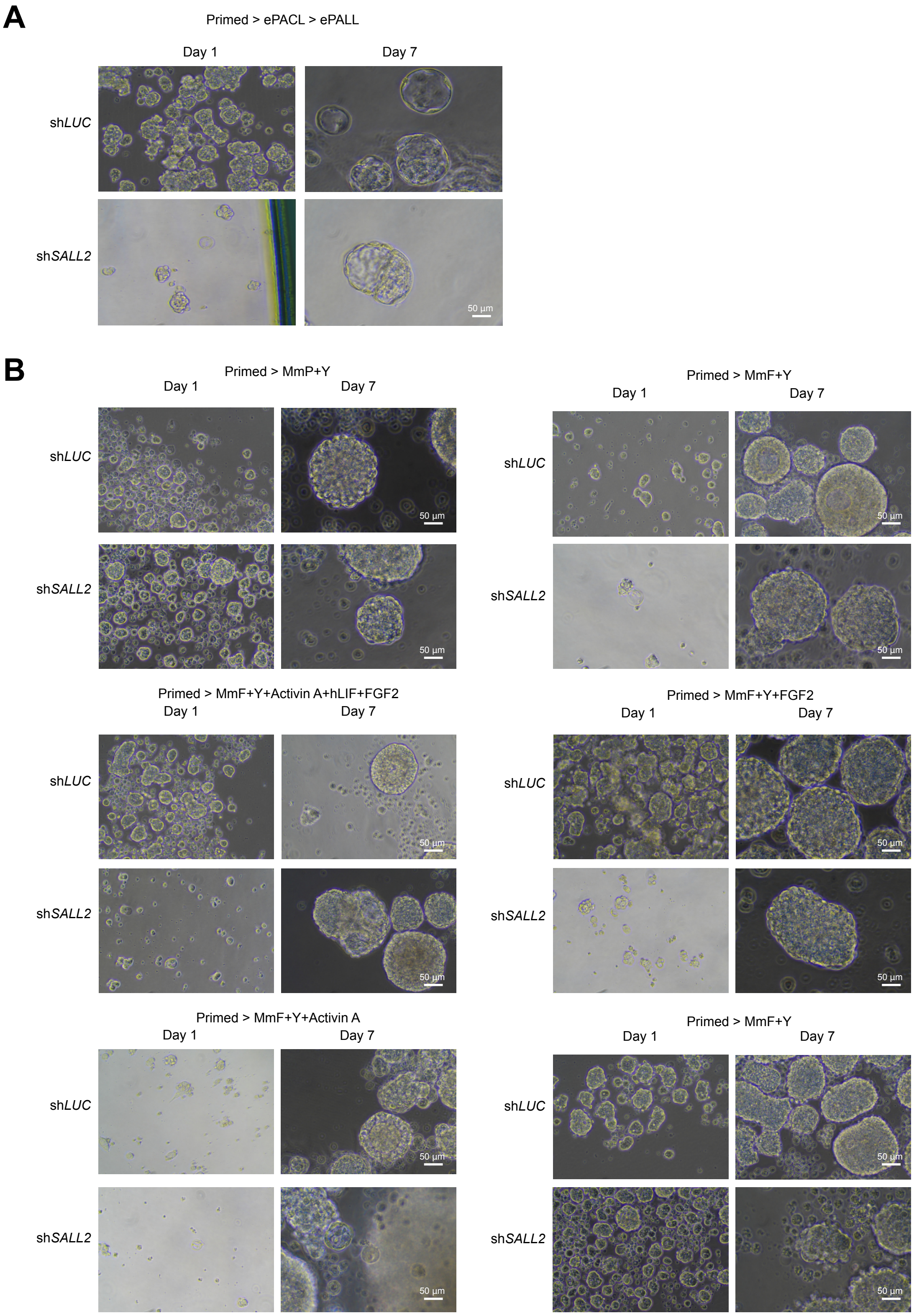


**Supplementary Figure 7**

**Supplementary Figure 7. The knockdown of *SALL2* affects blastoid formation.**

**A** Brightfield images of hPSC to blastoids grown in primed media, then ePACL, and ePALL, and transfected with the indicated shRNA. Scale bar = 50 um. missing part

**B** Brightfield images of hPSC to blastoids grown in primed media, then a minimal media omitting all inhibitors and growth factors, based on 5iLAF (MmF) or ePACL/PALL (MmP), plus the ROCK inhibitor Y-27632, or combinations of Activin A, LIF, FGF2, and transfected with the indicated shRNA. Scale bar = 50
